# Feedback loop involving AMPK, ERK and TFEB generates non-genetic heterogeneity in matrix-deprived cancer cells and regulates survival

**DOI:** 10.1101/736546

**Authors:** Saurav Kumar, Sayoni Maiti, Diptanshu Banerjee, Kishore Hari, Mohit Kumar Jolly, Annapoorni Rangarajan

## Abstract

Cancer cells metastasize by evading apoptosis induced due to the detachment from the extracellular matrix. Deciphering the adaptive strategies employed by cancer cells in matrix-deprived condition can help design novel therapeutic strategies to tackle metastasis. Here, we provide evidence for non-genetic heterogeneity in matrix-detached cells enabled by feedback loop among AMPK, ERK and TFEB that determines autophagy maturation and cell survival. The subpopulation of matrix-detached cells with pAMPK^low^/pERK^high^/TFEB^low^ state show autophagy maturation arrest and elevated cell death. Conversely, pAMPK^high^/pERK^low^/ TFEB^high^ cells show high autophagy maturation and better cell survival. We show that AMPK inhibits ERK activity in suspension; ERK negatively regulates TFEB which promotes autophagy maturation and re-enforces AMPK. Inhibition of ERK promotes autophagy maturation, cell survival and metastasis *in vivo*, while AMPK inhibition (and TFEB depletion) renders the population homogeneous by depleting the pAMPK^high^/pERK^low^/TFEB^high^ subpopulation, driving detachment-induced death. Such non-genetic heterogeneity is further deciphered by mathematical modelling and RNA-sequencing data of circulating tumor cells (CTCs) isolated from breast cancer patients. Altogether, our work unravels a contextual feedback loop involving two kinases and a transcription factor that helps a subpopulation to evade cell death in matrix-deprived condition. Disrupting such feedback loops may offer improved therapeutic efficacy and a novel approach to constrain metastasis.

**Significance of the Study:** Attachment to the extracellular matrix is pivotal for the growth and survival of normal epithelial cells. In contrast, cancer cells acquest the ability to survive matrix-deprivation and cause cancer spread, or metastasis, which is a leading cause of cancer-related deaths. Non-genetic heterogeneity within cancer cell populations is increasingly being recognized as a major cause of treatment failure. However, the implications of such heterogeneity in the survival of matrix-detached cancer cells remain poorly understood. In this study, we demonstrate feedback loops involving kinases and transcription factor to maintain non-genetic heterogeneous population, where population harbouring pAMPK^high^/pERK^low^/TFEB^high^ status shows survival advantage. Targeting such feedback loops that generate non-genetic heterogeneity can open newer therapeutic means to restrict cancer spread.

## Introduction

Normal epithelial cells need attachment to the extracellular matrix (ECM) for proper growth and differentiation. In contrast, detachment of cells from the ECM results in apoptosis, termed as “anoikis” [1, 2]. However, few cancer cells circumvent anoikis [3], survive during transit through the circulation, and subsequently seed metastasis – the major cause of cancer-related deaths. Therefore, understanding the adaptive mechanisms of cancer cells to deceive anoikis would help to restrain the spread of cancer.

Cancer cells evade anoikis by upregulating anti-apoptotic proteins such as Bcl-2, Bcl-XL, Bax, Bim and BAD, and activating pro-survival signaling which involves integrin, FAK, EGFR, TGFβ and PI3K [3–5]. Metabolic alteration is recently highlighted as a rapid adaptive mechanism of the cancer cells to survive in detachment [6, 7]. Previous work from our as well as other laboratories have elucidated the important role of AMP-activated protein kinase (AMPK), a cellular energy sensor and metabolic regulator, in providing anoikis resistance [8–10]. In addition, cellular processes such as epithelial-mesenchymal transition, stem-cell like properties, and entosis also aid in acquisition of anoikis resistance [4]. Yet, what mechanisms distinguish the cells that survive from the ones that undergo cell death remain to be understood.

Detachment of cells from the ECM induces autophagy, a cell survival mechanism in stress conditions [9, 11, 12]. Macroautophagy (or simply autophagy) is an evolutionarily conserved catabolic mechanism for degradation of protein aggregates, damaged organelles and intracellular pathogens within lysosomes [13]. It is a multistep-dynamic process that begins with the formation of a double membranous structure called autophagosome, which mature and fuse with lysosomes for the degradation of enclosed materials [13]. Cells consume these recycled key products from autophagy-mediated breakdown of cargo and survive several stresses including starvation, hypoxia, and anti-cancer therapeutics. In contrast, defective autophagy leads to neurological, cardiac and muscular pathologies [14]. Although several laboratories including ours have shown autophagy induction in response to matrix-detachment [9, 11, 12], but the fate of autophagy maturation is unknown. This is important because induction of autophagy but failure to complete the process can induce apoptosis [15].

Cancer cells show differential survival ability under matrix deprivation due to yet unknown mechanisms. Here, we demonstrate that matrix-deprived breast cancer cells show varying levels of ERK activity and autophagy maturation: the population of cells with high ERK activity show autophagy induction but maturation defect and high anoikis, whereas those with low ERK activity show autophagy maturation and better survival fitness. We identify a positive feedback loop involving AMPK, ERK and TFEB that enables non-genetic heterogeneity of cells with pAMPK^high^/pERK^low^/TFEB^high^ (high autophagy/low anoikis) and pAMPK^low^/ pERK^high^/TFEB^low^ (low autophagy/high anoikis) states. Inhibiting AMPK and TFEB depletion promotes elevation in ERK activity and impairs autophagy maturation, rendering cells unfit to survive in matrix-deprived conditions.

## Results

### Heterogeneous ERK signaling in matrix-deprived cells

In matrix-deprived cancer cells, both elevated as well as decreased ERK activity has been reported in different cell types [16–18]. These contrary reports led us to explore the role of ERK signaling in anoikis resistance of breast cancer cells. To do so, we reanalysed our previous transcriptomics data of MDA-MB-231 invasive breast cancer cells cultured in adherent (attached, Att) versus matrix-deprived (suspension, Sus) conditions for 24 hours [9]. We observed an increase in expression of genes related to ERK signaling in the suspension **(Figure S1A**). Additionally, GSEA analysis suggested enrichment of ERK signature genes in matrix-deprived condition **(Figure S1B)**. We further confirmed elevated ERK signaling by immunoblotting of whole cell lysate from multiple breast cancer cell lines (BT474, MDA-MB-231, MCF7) using phospho-ERK (Thr^202^/Tyr^204^)-specific antibodies **(Figure S1C)**. Levels of total ERK remained unaltered between these two conditions **(Figure S1C**). Further, we also observed an increase in *cFOS* expression **(Figure S1D)** and Egr1 promoter activity **(Figure S1E)** in suspension. Together, these results demonstrated elevation in ERK signaling in matrix-deprived breast cancer cells.

Cancer cells need to overcome anoikis not only while in circulation, but also until they find a proper niche at the secondary site. Therefore we checked the status of ERK signaling in long-term (one-week) cultures of cancer cells as spheroids in methylcellulose [8, 9]. Surprisingly, in contrast to cells subjected to matrix deprivation for 24 hours, anchorage-independent spheroids harvested after 7 days in culture revealed remarkably reduced levels of pERK **(Figure 1A)**. These contrasting observations of pERK status in single cell suspension versus spheroids led us to hypothesize the existence of heterogeneous ERK signaling in the matrix-deprived cells such that those cells with lower ERK activity are better fit to evade anoikis and generate anchorage-independent colonies.

**Figure 1.**
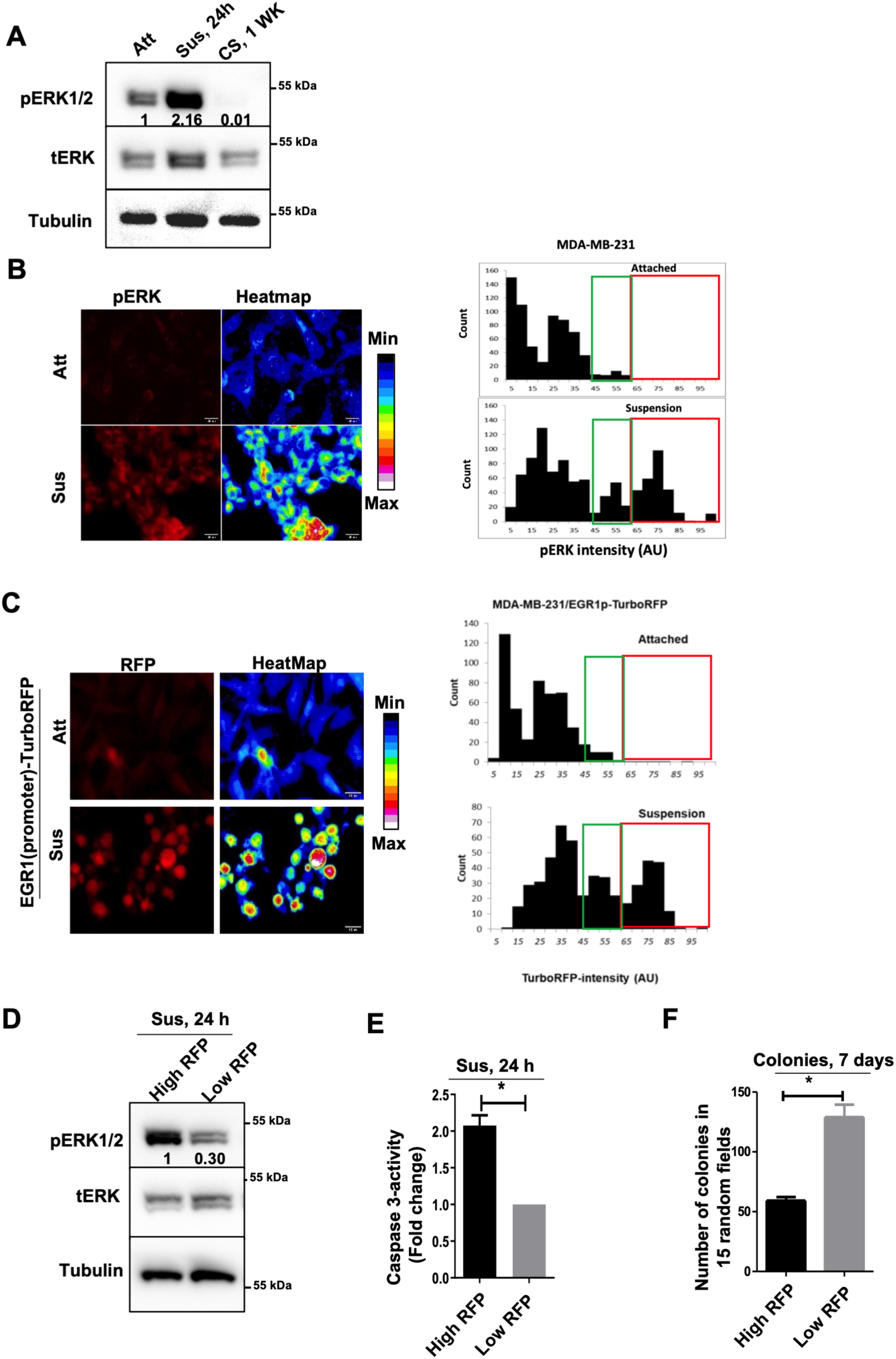
Heterogeneity in ERK activity in matrix-deprived conditions. A. Immunoblot analysis of BT-474 cells cultured in attached (Att), suspension condition (Sus) or anchorage independent cancer spheres (CS) in methylcellulose for 7-days; n=3. B. Fluorescent images of MDA-MB-231 cells cultured in attached (Att) or suspension (Sus) condition for 24 hours, probed for pERK, and visualized by confocal microscopy (Z-stack, scale bar, 20µM); n=3. Heat map was generated with the help of ImageJ. Histograms represent distribution of cells cultured in attached (Att) or suspension (Sus) conditions for 24 hours based on pERK intensity. C-F). MDA-MB-231 cells stably expressing EGR1_(promoter)_-TurboRFP were cultured in attached (Att) or suspension (Sus) conditions for 24 hours and imaged (C). High and low RFP subpopulations were separated by FACS sorting and harvested for immunoblotting (D); n=3, caspase-3 activity assay (E); n=3, and anchorage independent colonies formation (F); n=3. Error bars show mean and SEM from three biological repeats. *, P<0.05; Paired two tailed T-test.

To test this hypothesis, we first evaluated the status of pERK at individual cell level by immunostaining. Fluorescence microscopy revealed basal variation in pERK levels in MDA-MB-231 cells growing in adherent condition as unveiled by quantification of pERK intensities in individual cells (**Figure 1B**). In response to the matrix deprivation, we observed an overall increase in pERK levels with a more pronounced heterogeneity and the emergence of a subpopulation of pERK^high^ cells (**Figure 1B**, red box), (**Figure 1B**, histogram). We obtained similar data in several other cell lines, such as BT-474 (breast cancer) (**Figure S1F)**, HeLa (cervical cancer), OVCAR3 (ovarian cancer), and A549 (lung cancer) (**Figure S2**).

To access the significance of differential ERK activation under matrix-deprived condition, we employed the promoter reporter construct for ERK activity to isolate the high- and low- ERK activity cells. To do so, we established MDA-MB-231 cells stably expressing the Egr1-TurboRFP promoter-reporter system where the Egr1 promoter is driven by MEK/ERK signaling [19]. Fluorescence microscopy of Egr1-TurboRFP expressing cells yet again revealed a heterogeneous population of cells with variable ERK activity in matrix-detached cells compared to adherent cells (green and red boxes in **Figure 1C**). We isolated the low- and high-RFP cells using flow cytometer and confirmed higher pERK levels in high-RFP cells, and vice versa **(Figure 1D).** In accordance with our hypothesis, the low-RFP cells displayed less anoikis, as revealed by reduced caspase 3 activity **(Figure 1E),** and better colony formation potential, compared to the high-RFP cells **(Figure 1F)**. Altogether, these data revealed that matrix-detachment exacerbates differential ERK activity in a population of cells, where cells with low ERK activity have better survival fitness in matrix-deprivation conditions.

### ERK signaling negatively regulates autophagy maturation in matrix-deprived cells

We and others have previously shown that cancer cells adapt better to matrix-deprived condition by inducing autophagy [9, 11, 12]; yet the status of autophagy within individual cells remains unexplored. Therefore, we interrogated the status of autophagy in the matrix-deprived cells. To begin with, immunoblotting of whole cell population revealed increase in LC3-II levels in matrix-deprived MDA-MB-231 cells, which is suggestive of autophagy induction (**Figure S3A**). However, we simultaneously observed over 3-fold accumulation of p62/SQSTM1 in matrix-detached cells (**Figure S3A**). Elevated LC3-II along with p62 is indicative of autophagy maturation blockage [20]; hinting that autophagy may be induced but not completed by all cells in suspension.

Since ERK regulates fusion of autophagosomes with lysosomes, termed autophagy maturation [21–23], and matrix-deprived cells vary in ERK activity, therefore we hypothesized that cells in suspension also may have heterogeneity in autophagy maturation due to heterogenous nature of ERK signaling. To evaluate autophagy maturation on a cell-by-cell basis, we utilised the tandem-labelled mCherry-EGFP-LC3 reporter [24]. In this system, autophagosomes appear as yellow puncta (because of intact fluorescence from both mCherry and EGFP), whereas after fusion with lysosome, the autolysosomes are seen as red puncta (because of quenching of EGFP by the acidic nature of lysosome). The loss of EGFP fluorescence following autophagosome maturation results in increased mCherry/EGFP ratio which serves as a measure of autophagic flux. Firstly, we evaluated the behaviour of the mCherry-EGFP-LC3 reporter in adherent MDA MB 231 cells using an autophagy inducer rapamycin. As expected, treatment of rapamycin led to an increase in mCherry/EGFP ratio indicative of increased autophagic flux [25] **(Figure S3B).** Having confirmed the proper behaviour of mCherry-EGFP-LC3 reporter, we next subjected the cells to matrix deprivation. Interestingly, the distribution of mCherry/EGFP ratio from individual cells in suspension revealed two distinct populations of cells with the small, ∼25% of the cells showing elevated mCherry/EGFP ratio indicative of autophagosome acidification, while the vast majority showed equivalent signal from both RFP and EGFP indicative of poor acidification (**Figures 2A & S3C**).

**Figure 2.**
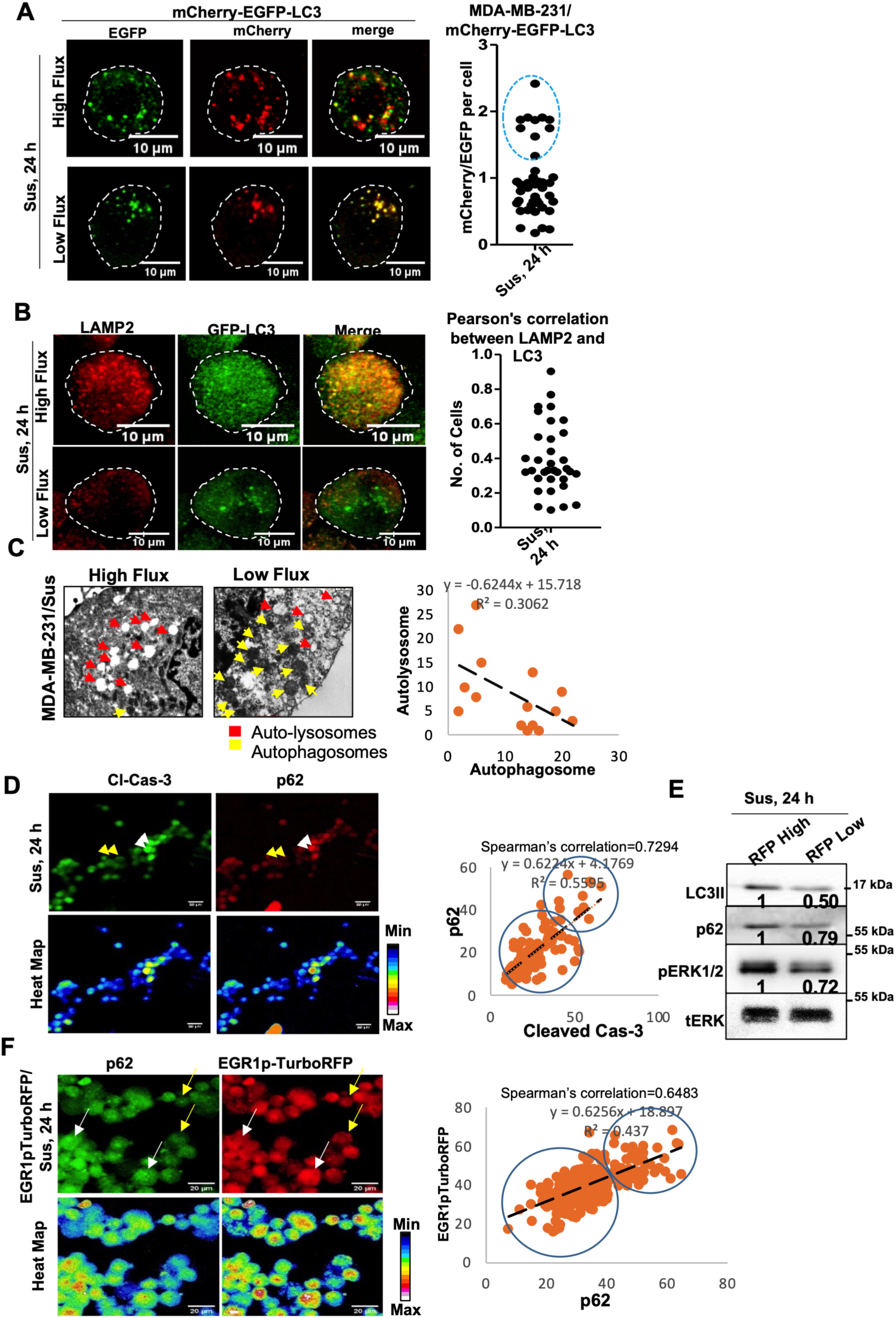
Matrix deprivation leads to heterogeneity in autophagy maturation. A. Fluorescent images of MDA-MB-231 cells stably expressing mCherry-EGFP-LC3 and cultured in suspension (Sus) conditions for 24 hours and visualized with confocal microscopy (Z-stack, scale bar, 10µM); n=3. Graph represents ratio of mChrrey/EGFP intensity. Circle indicates the cell population with high autophagy maturation. B. Immunofluorescence of MDA-MB-231 cells with anti-LAMP2 antibody (Cy3) and co-localization with GFP-LC3 puncta in cells cultured in suspension (Sus) conditions for 24 hours and visualized with confocal microscopy (Z-stack, scale bar, 10 µM); n=3. Colocalization of LAMP2 and LC3 was measured by Pearson’s correlation coefficient employing Coloc-2 plugin in ImageJ and represented as dot plot (each dot represents single cell). C. Electron micrographs of MDA-MB-231 cells cultured in matrix-deprived conditions for 24 hours. Yellow arrows denote autophagosomes; red arrows indicate autolysosomes. Quantification of autophagosomes and autolysomes from electron micrographs (*n* = 15 fields). D. Immunofluorescence of MDA-MB-231 cells with anti-cleaved-caspase-3 or anti-p62 antibody cultured in suspension (Sus) condition for 24 hours (yellow arrow represents less cleaved caspase-3^less^/p62^less^ and white arrow represents cleaved caspase 3^high^/p62^high^ level). Heat map was generated with the help of ImageJ. Spearman correlation analysis between cleaved-caspase-3 and p62 was plotted using excel. E. MDA-MB-231 cells stably expressing EGR1_(promoter)_-TurboRFP were cultured in suspension (Sus) conditions for 24 hours. High and low RFP subpopulations were separated by FACS sorting and harvested for immunoblotting; n=3. F. Immunofluorescence of MDA-MB-231 cells stably expressing EGR1_(promoter)_-TurboRFP with anti-p62 antibody cultured in suspension (Sus) conditions for 24 hours (yellow arrow represents RFP^less^/p62^less^ and white arrow represents RFP^high^/p62^high^). Heat map was generated with the help of ImageJ. Spearman correlation analysis between p62 and TurboRFP was plotted using excel.

To further assess the status of autophagy maturation we analysed the co-localization of LC3 with LAMP2, a lysosomal marker [26]. Large number of MDA-MB-231 cells stably expressing GFP-LC3 displayed less co-localization with LAMP2 (70%) in matrix-deprivation, thus validating defective autophagy maturation **(Figures 2B & S3D)**. Electron microscopy further established this data demonstrating an inverse correlation between numbers of autophagosomes and autolysosomes within individual cells (**Figure 2C**). These data revealed heterogeneity in autophagy maturation in matrix-detached cells with a large number (∼75%) of cells showing defective autophagy maturation.

We further investigated the relationship between autophagy maturation and anoikis. To do so, we first compared the level of cleaved caspase 3 (a marker of apoptosis) with that of accumulated p62 (a marker of blocked autophagy) in individual cells by immunocytochemistry (**Figure 2D**). A Spearman’s correlation analysis revealed a positive correlation between cleaved caspase 3 and p62 levels (**Figure 2E**), suggestive of elevated anoikis in cells with impaired autophagy maturation. Conversely, we observed less accumulation of p62 in anchorage-independent colonies **(Figure S4A)**.

So far, our data revealed the existence of heterogeneity in ERK signaling as well as autophagy maturation. We next addressed the relationship between ERK activity and autophagy maturation. To do so, we investigated the status of autophagy in the FACS sorted low-RFP (ERK^low^) and high-RFP (ERK^high^) cells. Compared to ERK^high^ cells, immunoblotting revealed lesser LC3-II as well as p62 levels in ERK^low^ cells, suggesting high autophagy maturation in cells with low ERK activity **(Figure 2F**). To further confirm this negative correlation between ERK signaling and autophagy maturation, we investigated the colocalization of endogenous LC3 (green) and LAMP2 (magenta) in the EGR1-turboRFP promoter-reporter expressing stable cells. Quantitative confocal microscopy revealed a higher co-localization of LC3 and LAMP2 in cells with low RFP expression (**Figure S4B)**, thus implying higher autophagy in cells with low ERK activity. Immunostaining for p62 in these cells further supported this data as it showed a positive corelation between RFP intensity and p62 accumulation **(Figure 2G)**.

Conversely, immunostaining for pERK in mCherry-EGFP-LC3 expressing cells revealed higher pERK in cells with low autophagy flux (yellow arrows) and vice-versa (**Figure S4C**). Taken together, these data suggest a differential ERK signaling and autophagy maturation in matrix-detached cells: those with low ERK activity have elevated autophagy maturation and show better survival.

### AMPK-mediated phosphorylation of PEA15 inhibits MEK-ERK interaction and activity

We next sought to understand the regulators of ERK activity in matrix-detached cells. Work done in our laboratory and that of others has identified AMPK activation in response to matrix-deprivation [8, 27]. To investigate if AMPK is involved in regulating ERK activity in suspension, we first measured AMPK activity in RFP-high and RFP-low cells by measuring the levels of its phosphorylated bonafide substrate, ACC. We observed an inverse corelation between AMPK and ERK activities such that RFP-high cells showed less pACC levels and vice versa, while total protein levels of ACC and ERK remain unchanged **(Figure 3A & S5A)**. Quantitative immunofluorescence analysis of pACC and pERK levels in matrix-detached MDA-MB-231 cells further confirmed this inverse relationship between AMPK activity and pERK levels **(Figure 3B)**. To further elucidate this inverse co-relation in *in vivo*, we resorted to the mouse lactating female mammary gland [12]. The matrix-deprived luminal cells of the lactating mammary gland also showed inverse correlation between pAMPK and pERK status **(Figure 3C)**. Importantly, RNA-sequencing data of CTCs isolated from breast cancer patients revealed inverse corelation in AMPK and ERK gene signature (**Figure 3D**). Together these results suggest a negative corelation between AMPK and ERK signaling.

**Figure 3.**
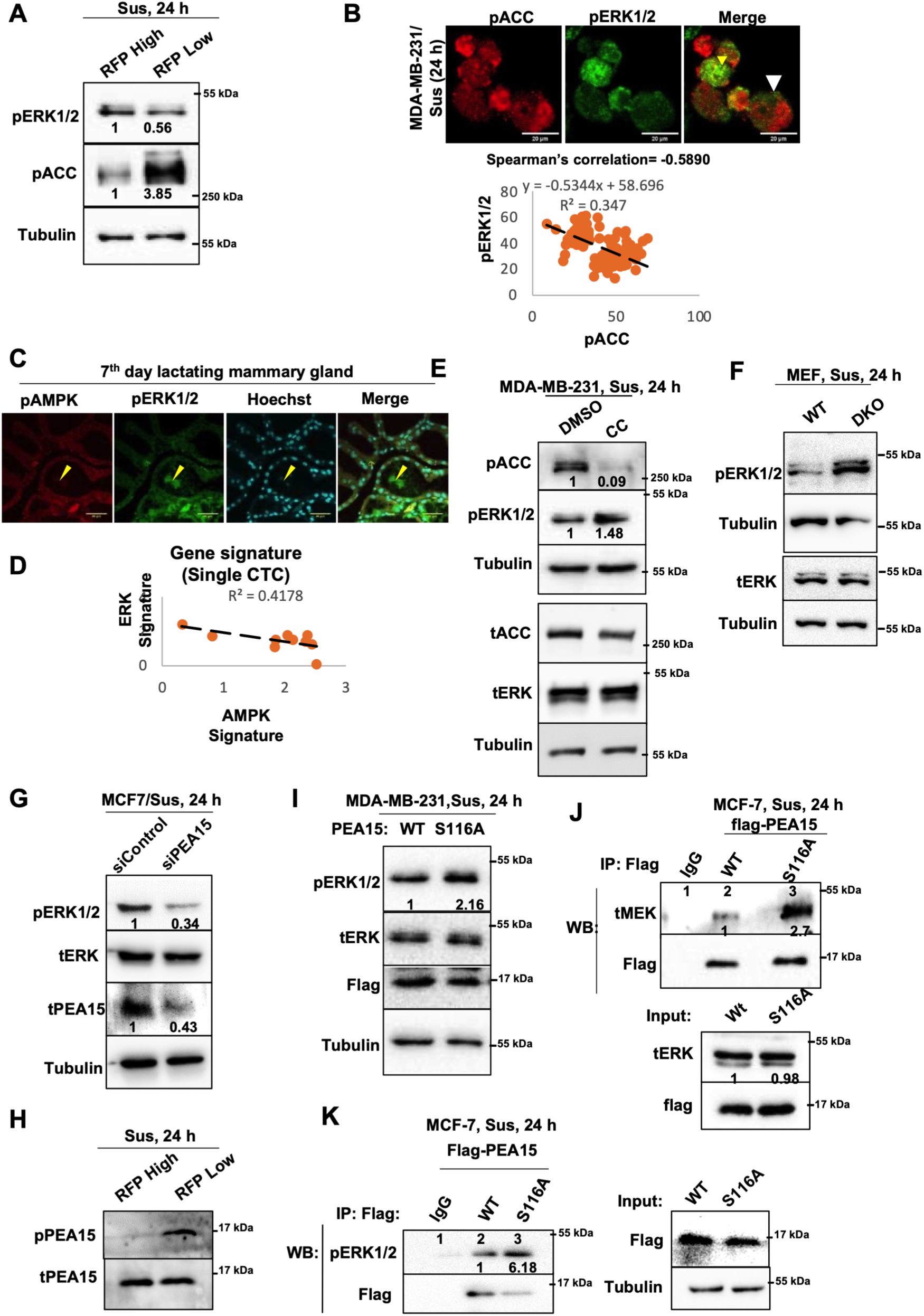
AMPK inhibits ERK activity via PEA15 and promotes heterogeneity. A. MDA-MB-231 cells stably expressing EGR1_(promoter)_-TurboRFP were cultured in suspension (Sus) condition for 24 hours. High and low RFP subpopulations were separated by FACS sorting and harvested for immunoblotting; n=3. B. Immunofluorescence of MDA-MB-231 cells with anti-pACC or anti-pERK antibody cultured in suspension (Sus) condition for 24 hours (yellow arrow represents pACC^low^/pERK^high^ and white arrow represents pACC^high^/pERK^low^); n=2. Co-relation graph was plotted using excel. Spearman correlation analysis between pACC and pERK was plotted using excel. C. Immunofluorescence of lactating mammary glands of mouse for anti-pAMPK and anti-pERK and visualised with confocal microscopy (Z-stack, scale bar, 40 µM); n=3. D. Double-label immunofluorescent staining of pACC or pERK in frozen sections of primary breast cancer patient. Blue dotted areas represent cells with pERK^high^/pACC^low^ and white dotted area represents cells with pERK^low^/pACC^high^; n=4. E. Scatter and co-relation graph was plotted for the expression of ERK and AMPK dependent genes from publicly available RNA-sequencing data for 15 single CTCs from ten breast cancer patients (<u>GSE51827</u>). F & G). Immunoblots analysis of following cell lysates were harvested and probed for specified proteins: (F). MDA-MB-231 cells were cultured in suspension (Sus) condition for 24 hours in presence of vehicle control (DMSO) or AMPK inhibitor (compound C, CC); n=3. (G). Wild type (WT) and AMPKα1/α2-double knockout (DKO) mouse embryonic fibroblast (MEF) cultured in suspension (Sus) for 24 hours; n=2. (H). MCF7 cells cultured in suspension (Sus) for 24 hours after treated with control or PEA15 siRNA; n=2. (I). MDA-MB-231 cells stably expressing EGR1_(promoter)_-TurboRFP were cultured in suspension (Sus) condition for 24 hours. High and low RFP subpopulations were separated by FACS sorting; n=3. (J). MDA-MB-231 cells stably overexpressing Flag-tag wild type PEA15 (WT-PEA15) or nonphosphorylatable mutant of PEA15 (S116A-PEA15) cultured in suspension (Sus) condition for 24 hours; n=5. K & L). Immunoblot analysis of immunoprecipitated (IP) products with IgG control or anti-tPEA15 antibodies from MCF-7 cells transiently transfected with Flag-tag WT-PEA15 or S116A-PEA15 cultured in suspension (Sus) condition. 2% of the whole-cell lysate was used as input and probed for specified proteins; n=3.

To further address the role of AMPK in regulating pERK levels in suspension, we interrogated the effect of pharmacological inhibition as well as genetic ablation of AMPK on pERK levels. Treatment of matrix-deprived MDA-MB-231 cells with a pharmacological inhibitor of AMPK compound C (henceforth referred to as CC) [28] led to a reduction in pACC levels, confirming the efficacy of the inhibitor **(Figure 3E).** Simultaneously, we observed an increase in pERK levels compared to vehicle DMSO treated cells **(Figure 3F)**; while CC had no effect on total ERK levels **(Figure 3F)**. We observed similar effects of AMPK inhibition in yet another breast cancer cell line BT-474 **(Figure S5B)**. We further confirmed this finding in the double knockout for AMPK α1/α2 (MEF-DKO) cells in suspension **(Figure 3G).** Inversely, we did not observe decrease in ERK activity upon activating AMPK in the attached conditions upon A76 treatment (**Figure S5C**). Altogether, leveraging pharmacological as well as genetic approaches to impair AMPK, we uncovered that AMPK inhibits ERK activity in matrix-deprived condition.

We next investigated the mechanism underlying AMPK-mediated negative regulation of ERK activity in suspension. Levels of phosphorylated and active form of MEK, upstream kinase of ERK, did not change in response to matrix deprivation or AMPK inhibition (**Figures S5D and S5E, respectively**). Next, we asked if AMPK affects MEK-ERK interaction. We had previously identified phosphoprotein enriched in astrocytes 15 kDa (PEA15) as a substrate of AMPK in suspension leading to S116 phosphorylation [8]. PEA15 regulates ERK activity by binding and preventing its nuclear entry [29], however, its role in regulating ERK phosphorylation is unknown. Therefore, we depleted PEA15 and looked for pERK status in matrix-deprived cells. Interestingly, PEA15 depletion led to decrease in pERK levels with unaltered tERK1/2 (**Figure 3G**). Next, we investigated the role of AMPK-mediated S116 phosphorylation of PEA15 in impairing ERK phosphorylation. We observed higher levels of phospho S116-PEA15 in RFP-low cells and vice versa **(Figure 3H)**. Additionally, overexpression of non-phosphorylatable S116A mutant of PEA15 led to an increase in pERK levels in matrix-deprived condition (**Figure 3I**), but not in attached cells (**Figure S5F**). Moreover, treatment with AMPK inhibitor Compound C (CC) resulted in elevated pERK levels in WT-PEA15 expressing cells, but not in S116A-PEA15 expressing cells, in suspension **(Figure S5G)**. Together, these data uncovered a contextual role of serine 116 phosphorylated PEA15 in the negative regulation of ERK signaling in matrix-deprived condition.

Given that MEK-ERK interaction is important for the phosphorylation and activation of ERK, therefore, we checked if pS116-PEA15 inhibits MEK-ERK interaction. To address this, we immunoprecipitated PEA15 from cells expressing flag-tagged WT or S116A-PEA15 mutant using anti-flag antibody. Immunoprecipitation with flag antibody followed by immunoblotting revealed presence of MEK and ERK in the complex (**Figures 3J and K**, respectively). Interestingly, we observed higher levels of MEK in S116A-PEA15 expressing cells compared to WT-PEA15 (**Figure 3J).** Consistent with this, we also detected elevated levels of pERK in S116A-PEA15 expressing cells **(Figure 3K)**. Further, immunoprecipitation of endogenous PEA15 revealed elevated level of MEK and pERK upon AMPK inhibition **(Figure S5H**). Altogether, these data suggest a novel role for AMPK-PEA15 axis in negative regulation of ERK phosphorylation in matrix-deprived condition.

### ERK regulates autophagy by negatively regulating TFEB levels in suspension

After deciphering AMPK-PEA15 axis upstream to ERK in suspension, we focussed on downstream signaling that contributes to autophagy maturation and anoikis resistance. Transcription factor EB (TFEB) is a master regulator of lysosomal biogenesis and autophagy [30]. Immunoblotting of the Egr1-Turbo-RFP promoter reporter activity-based FACS-sorted cells showed considerably high levels of TFEB in low-RFP cells as compared to the counter cells (**Figure 4A**). Likewise, immunofluorescence for TFEB in the unsorted population expressing the Egr1-Turbo-RFP promoter reporter revealed an inverse correlation between RFP and TFEB levels (**Figure S6A)**, together suggesting that ERK signaling may negatively regulate TFEB level. Consistent with this, ERK inhibition led to an increase in TFEB levels in suspension (**Figures 4B & S6B)**, but not that of TFE3 (**Figure S6C**), another related protein that regulates lysosomal functions. Further AMPK inhibition led to reduction in TFEB levels (**Figure 4C**), while simultaneous ERK inhibition reverted this (**Figure 4D**), implying the involvement of AMPK-ERK axis in the regulation of TFEB levels in suspension.

**Figure 4.**
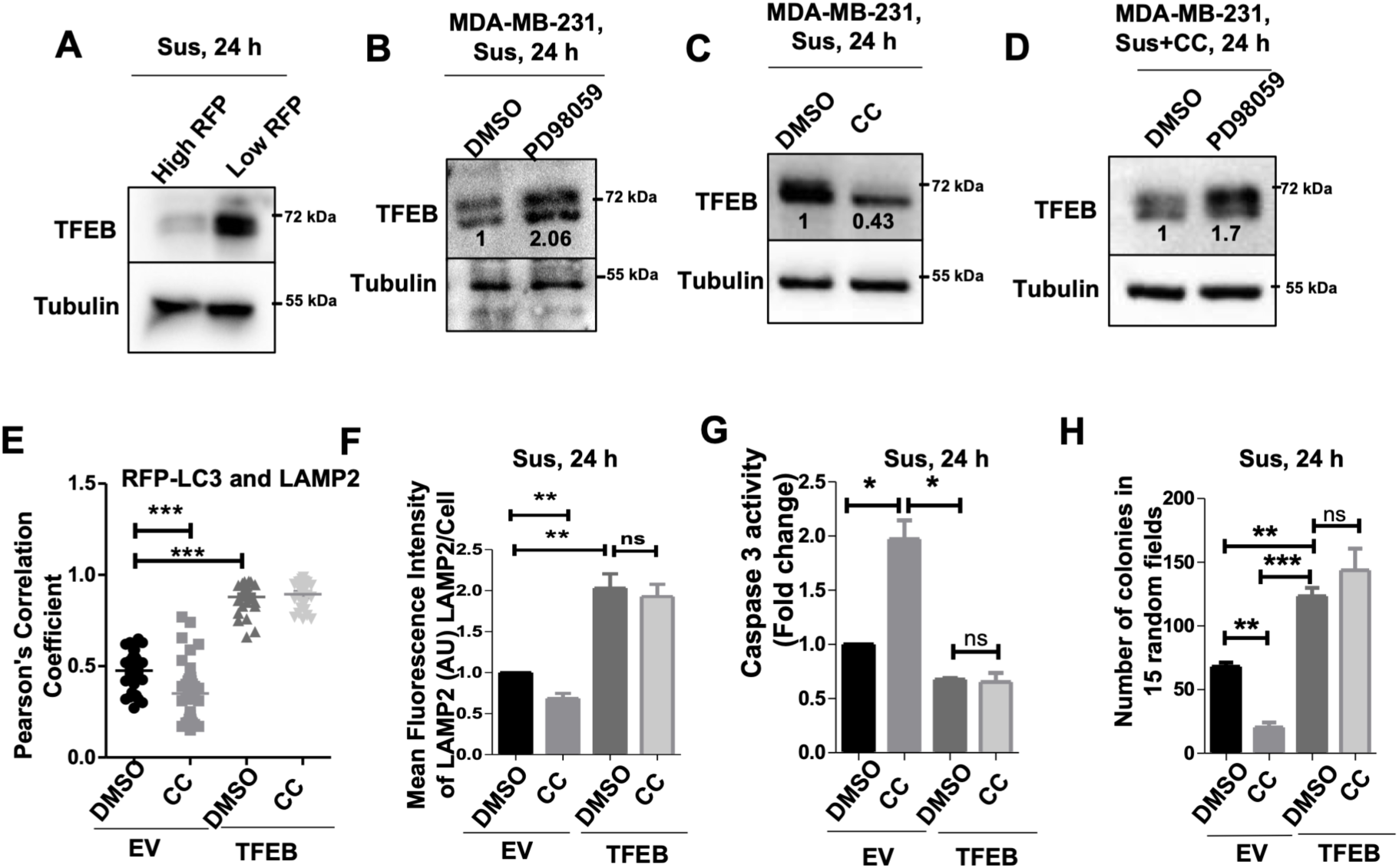
AMPK upregulates TFEB level by inhibition of ERK activity in matrix-deprived condition. A-D). Immunoblots of following cell lysates were harvested and probed for specified proteins: (A). MDA-MB-231 cells stably expressing EGR1_(promoter)_-TurboRFP were cultured in suspension (Sus) condition for 24 hours. High and low RFP subpopulations were separated by FACS sorting; n=3. (B). MDA-MB-231 cells cultured in suspension (Sus) for 24 hours in presence of vehicle control (DMSO) or MEK-inhibitor (PD98059); n=3. (C). MDA-MB-231 cells cultured in suspension for 24 hours in presence of vehicle control (DMSO) or AMPK inhibitor (CC); n=3. (D). MDA-MB-231 cells cultured in suspension for 24 hours in presence of AMPK inhibitor (compound C) plus treated with vehicle control (DMSO) or MEK-inhibitor (PD98059); n=3. E. Dot plot showing Pearson’s coefficient as a measure of colocalization of LAMP2 and RFP-LC3 in MDA-MB-231 cells stably overexpressing control empty vector or EGFP-TFEB were cultured in suspension (Sus) condition for 24 hours in presence of vehicle control (DMSO) or AMPK inhibitor (CC) (also see **S8B**); n=3. F. Graph represents mean fluorescence intensity of LAMP2 in MDA-MB-231 cells stably overexpressing control empty vector or EGFP-TFEB cultured in suspension (Sus) conditions for 24 hours in presence of vehicle control (DMSO) or AMPK inhibitor (compound C) (≥50 cells analysed per sample/experiment) (also see **S8C**); n=5. G & H). MDA-MB-231 cells stably expressing control empty vector and EGFP tagged TFEB cultured in suspension (Sus) conditions for 24 hours in presence of vehicle control (DMSO) or AMPK inhibitor (compound C) and harvested for flow cytometric analysis of caspase-3 activity (G) and anchorage independent colonies formation (H); n=2.

ERK inhibition in suspension also led to an increase in exogenously expressed EGFP-TFEB levels (**Figure S7A**), which might suggest a regulation of TFEB stability by ERK. Consistent with this, cycloheximide chase assay revealed a delayed decrease in TFEB protein level in suspension upon ERK inhibition by PD98059 compared to DMSO treated cells (**Figure S7B)**. Mass spectrometry-based analysis of TFEB-interactome unravelled its interaction with SUMO3 protein that was enriched after AMPK inhibition, implying a possible regulation of TFEB degradation in suspension through AMPK-ERK axis **(Figure S7C)**.

We further examined the role of TFEB downstream of AMPK-ERK axis in regulating autophagy maturation and anoikis. Since high ERK activity correlates with reduced TFEB levels, we asked if TFEB overexpression can enhance autophagy maturation and anoikis resistance. Overexpression of EGFP-TFEB led to increase in LAMP2 levels in matrix-deprived cells, as visualized by immunofluorescence microscopy **(Figure S8A).** Importantly, quantitative confocal microscopy revealed increased co-localisation of LAMP2 with LC3-RFP in TFEB overexpressing cells **(Figures 4E & S8B)**, suggesting increased autophagy maturation. Furthermore, TFEB overexpression reverted AMPK inhibition-mediated decrease in LAMP2 levels **(Figures 4F & S8C)**, autophagic-maturation defect **(Figures 4E & S9A)** and anoikis **(Figure 4G)**. Reinforcing this data, TFEB overexpression led to a significant increase in the number of anchorage-independent colonies formed, as well as rescued the colony formation deficiency imposed by AMPK inhibition **(Figure 4H)**. Altogether, these experiments emphasize that TFEB upregulation downstream of AMPK-ERK in suspension promotes autophagy maturation to overcome anoikis and survive better.

### Bi-directional positive feedback loop involving AMPK, ERK and TFEB in suspension

Our data thus far revealed at least two subpopulations in suspension **–** pAMPK^high^/pERK^low^/TFEB^high^ and pAMPK^low^/ pERK^high^/TFEB^low^ **(Figures 1D, 3 & 4A)**, demonstrating systems with two cell states (bistability). Bistability usually emerges in cases of ‘double positive’ or ‘double negative’ feedback loops between a set of molecular factors [31]. To examine this possibility in suspension, we investigated possible cross talks between TFEB and AMPK/ERK signaling. We have demonstrated that AMPK-ERK axis regulates TFEB levels; we now asked if TFEB in turn regulates AMPK and ERK signaling. Interestingly, in EGFP-TFEB overexpressing matrix-deprived MDA-MB-231 cells, immunoblotting revealed increase in pAMPK and pACC levels, suggestive of elevated AMPK activity **(Figures 5A)**. We observed elevated pACC levels in yet another cell line MCF7 overexpressing EGFP-TFEB **(Figure S9A).** Additionally, we also observed reduction in pERK levels upon TFEB overexpression **(Figures 5A & S9A**). Thus, overexpression of TFEB leads to hyperactivation of AMPK and inhibition of ERK activity **(Figure 5A)**, while we previously saw ERK negatively regulates TFEB levels **(Figures 4B & 4D)**; together these data suggest possible feedback loops involving AMPK, ERK, and TFEB.

**Figure 5.**
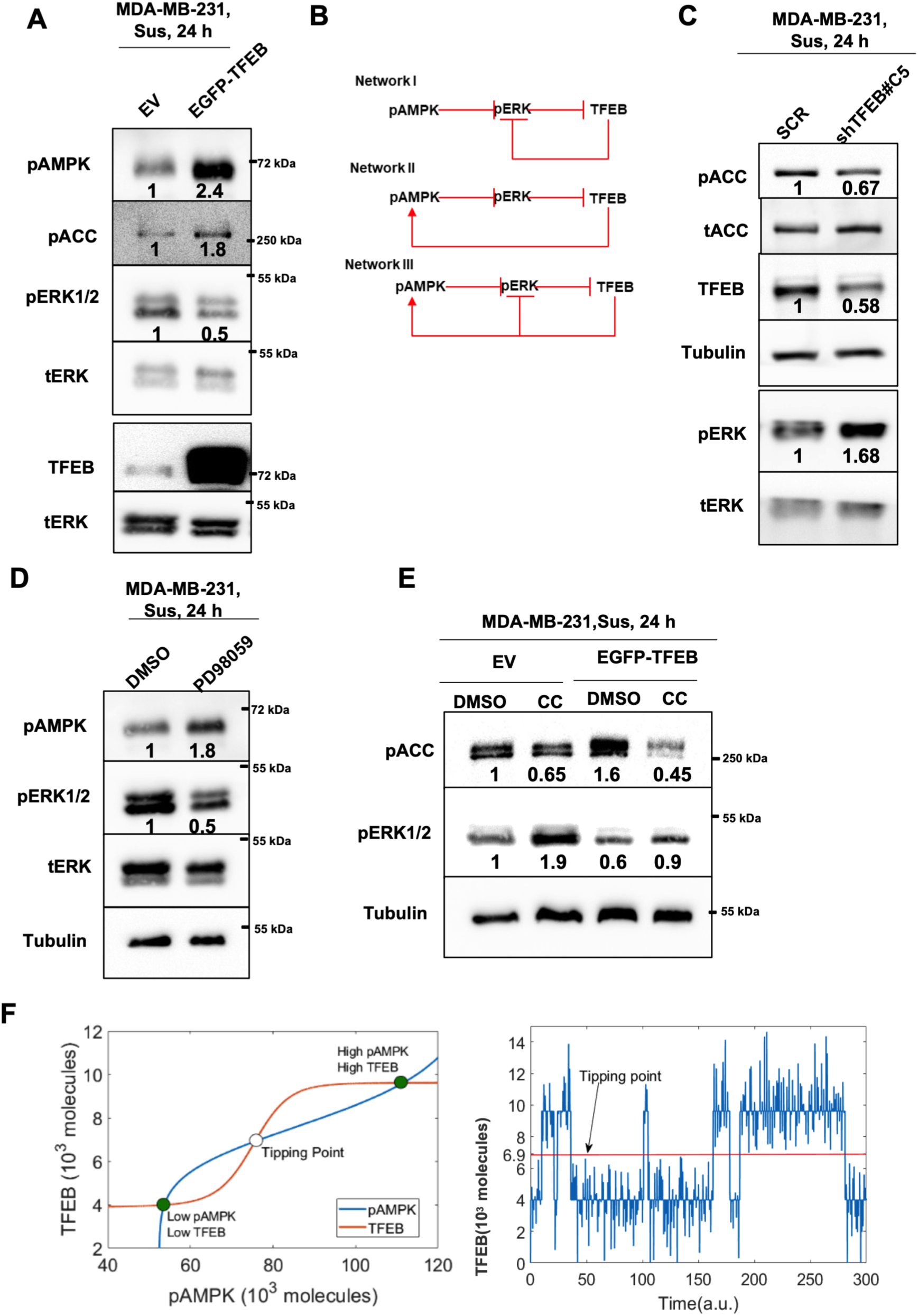
TFEB-mediated upregulation of AMPK activity. A. Immunoblot analysis of MDA-MB-231 cells stably overexpressing control empty vector (EV) or EGFP-TFEB were cultured in suspension (Sus) condition for 24 hours; n=3. B. Potential feedback loops that can represent possible cross talk between AMPK, ERK, and TFEB in matrix-deprived condition. C. MDA-MB-231 cells stably overexpressing Scr control or shTFEB#C5 cultured in suspension (Sus) condition for 24 hours and harvested for immunoblotting; n=3. D & E). Immunoblots analysis of (D). MDA-MB-231 cultured in suspension (Sus) condition for 24 hours in presence of vehicle control (DMSO) or MEK-inhibitor (PD98059) for 24 hours; n=3. (E). MDA-MB-231 cells stably overexpressing control empty vector or EGFP-TFEB were cultured in suspension (Sus) condition for 24 hours in presence of vehicle control (DMSO) or AMPK inhibitor (CC). n=3. F. (left) Nullclines generated by the mathematical model for network II; they represent the change of steady state concentration of pAMPK with change in TFEB concentration (blue) and vice versa (red). The intersections of the two curves represent the steady states of the system. Intersections highlighted in green are stable steady states, i.e., cellular phenotypes, while that highlighted in white represents the “tipping point” for transitions among these phenotypes. (right) Stochastic variations can lead to cells dynamically switching among the two phenotypes upon various perturbations, once they cross the tipping point (highlighted by red line).

We tested for three different potential feedback loops: **I)** TFEB inhibits ERK without involving AMPK (a ‘double negative’ one between ERK and TFEB), **II)** TFEB activates AMPK directly or indirectly (a ‘double positive’ one between AMPK and TFEB), and **III)** a combination of both of the above-mentioned possibilities **(Figure 5B)**. To examine which of these feedback loops actually operates within matrix-detached cells, we checked the effects of TFEB depletion, ERK inhibition, and AMPK inhibition. TFEB depletion resulted in reduced pACC levels **(Figures 5C and S9B)** and elevated pERK levels **(Figure 5C)**. ERK inhibition led to elevated pACC levels **(Figure 5D),** and we have previously shown that ERK inhibition led to elevated TFEB levels (**Figure 4D**). Moreover, we observed decrease in pACC levels in cells stably expressing S116A-PEA15 **(Figure S9C)** where we had previously observed higher pERK levels **(Figure 3H).** We next tested the effect of AMPK inhibition in TFEB-overexpressing cells and measured the levels of pERK. We observed no change in pERK level upon inhibition of AMPK activity in matrix-deprived cells over-expressing TFEB (**Figure 5E**, compare lane 3 and 4^th^), while control cells treated with AMPK-inhibitor displayed increased pERK activity (**Figure 5E**, compare lane 1^st^ and 2^nd^). This observation can be supported by network II, but not by network I & III (**Figure 5B)**. According to network III, the above-mentioned experiment **(Figure 5E)** would have led to a decrease in pERK levels, which we did not observe. Therefore, TFEB may activate AMPK through as yet unidentified players leading to the formation of a ‘double positive’ feedback loop between pAMPK and TFEB, and consequently a decreased ERK activity **(network II) (Figure 5B & 5E)**.

We constructed a mathematical model capturing the interactions shown in network II. This network can give rise to two distinct cellular phenotypes – pAMPK^high^/TFEB^high^/pERK^low^ and pAMPK^low^/ TFEB^low^/pERK^high^ (shown by solid green circles in **Figures 5F & S9D**) as observed in our experiments. Our mathematical model predicts that cells in these two states can also switch spontaneously, under sufficiently strong stochastic/noise perturbations, once they cross a ‘tipping point’ (shown by white circles in **Figures 5F & S9D**). This prediction is largely robust to parameter variation in the model (**Figure S9E**) and consistent with our experimental observations of a switch in phenotype upon overexpression or inhibition of TFEB (**Figure 5**).

### Disrupting the AMPK-ERK-TFEB feedback loop promotes anoikis

Having identified a feedback loop involving AMPK-ERK-TFEB in suspension, we investigated the biological relevance of this by testing the effects of inhibition/depletion of ERK, AMPK and TFEB on autophagy maturation and anoikis.

We reasoned that treatment with ERK inhibitor would abolish pERK heterogeneity in the whole cell population, and consequently improve overall survival under matrix deprivation. Treatment of matrix-deprived cells with MEK inhibitor PD98059 resulted in reduced pERK levels (**Figure S10A**). Additionally, we observed an increase in autophagy maturation in these cells as suggested by a decrease in the levels of LC3-II together with p62 **(Figure S10A)**. Similar results were confirmed in yet another breast cancer cell line, BT-474 **(Figure S10B)**. We further used a genetic approach to reduce pERK levels by overexpressing dominant-negative form of MEK1 (MEK1-K97A). We observed a decrease in the levels of pERK, LC3-II and p62 in these cells **(Figure S10C)**. These data suggest that inhibition of ERK activity can elevate autophagic flux at the whole population level.

We further examined the effects of ERK inhibition on autophagy maturation in individual cells. Treatment with PD98059 also led to an increase in red punctae in a large number of matrix-deprived cells expressing mCherry-GFP-LC3 **(Figure S10D)**. Furthermore, we observed a shift towards increased co-localization of LAMP2 with GFP-LC3 upon ERK inhibition **(Figure 6A)**. Consistent with this, we observed a reduction in anoikis upon ERK inhibition, as revealed by reduced caspase 3 activity (**Figure S10E)**, and increase in the number of anchorage independent colonies **(Figures S10F & 6B**). These data together suggest that reduction in ERK activity in matrix deprived cells promotes autophagy maturation and anoikis resistance.

**Figure 6.**
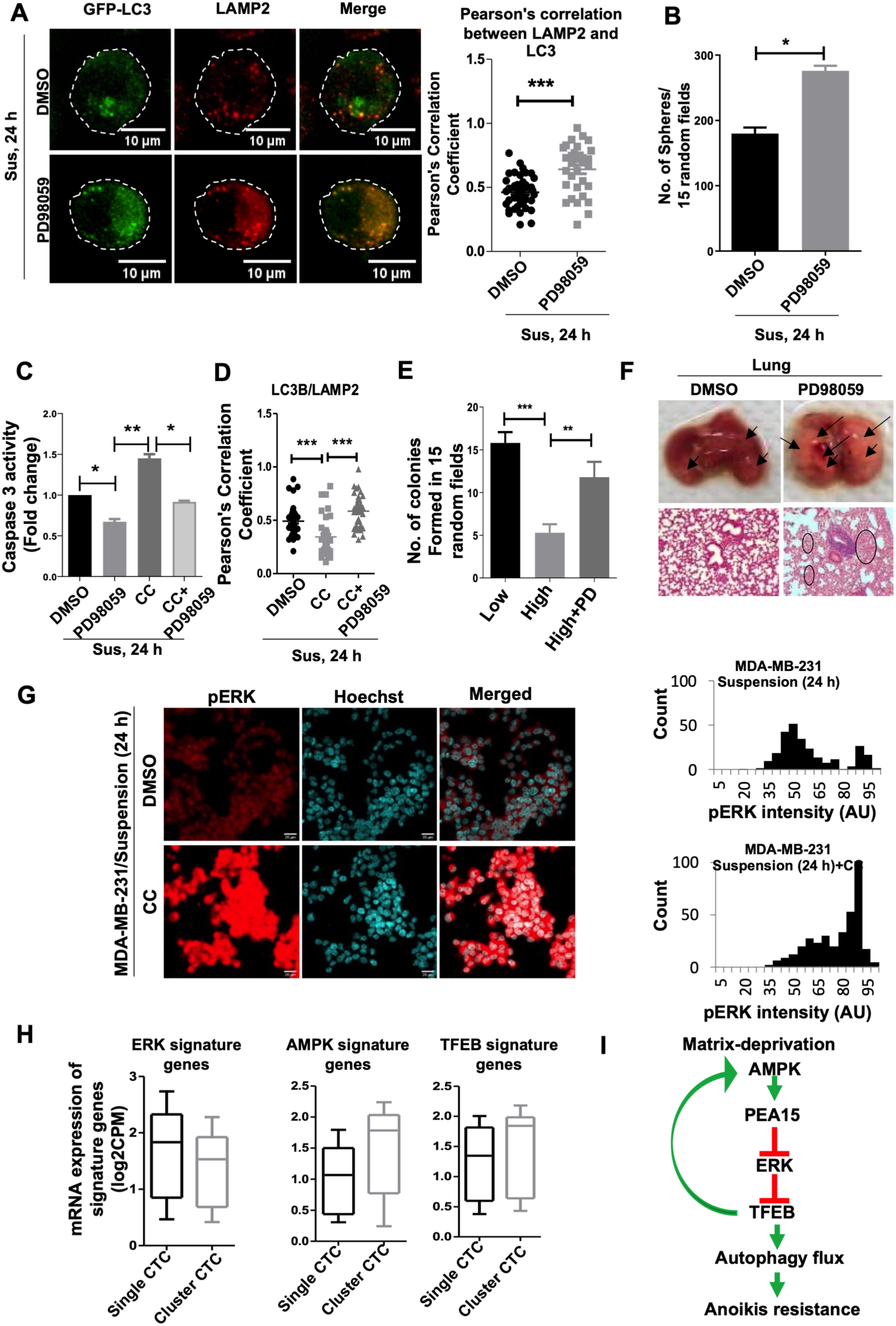
Breaking the AMPK-ERK feedback loop promotes anoikis in breast cancer cells. A. Immunofluorescence of MDA-MB-231 cells with anti-LAMP2 antibody (Cy3) and co-localization with GFP-LC3 puncta in cells cultured in suspension (Sus) condition for 24 hours in presence of DMSO or PD98059 and visualized with confocal microscopy (Z-stack, scale bar, 10 µM); n=3. Colocalization of LAMP2 and LC3 was measured by Person’s correlation coefficient employing Coloc-2 plugin in ImageJ and represented as dot plot (each dot represents single cell). B. Graph represents number of anchorage independent colonies formation from MDA-MB-231 cells cultured in presence of vehicles control (DMSO) or PD98059; n=3. C. Flow cytometric analysis of caspase-3 activity for MDA-MB-231 cells cultured in suspension (Sus) condition for 24 hours in presence of vehicle control (DMSO), MEK-inhibitor (PD98059), AMPK inhibitor (compound C, CC) with or without MEK-inhibitor (PD98059) for 24 hours; n=3. D. Dot plot showing Pearson’s coefficient as a measure of colocalization of LAMP2 and GFP-LC3 in MDA-MB-231 cells cultured in suspension (Sus) condition for 24 hours in presence of vehicle control (DMSO) or AMPK inhibitor (compound C, CC) with or without MEK-inhibitor (PD98059) (also see **Figure S11A**); n=3. E. Graph represents number of anchorage independent colonies formation from MDA-MB-231 cells stably expressing EGR1_(promoter)_-TurboRFP sorted for high and low RFP, where high RFP cells were cultured in the presence of PD98059; n=3. F. Representative images of lung metastasis following intraperitoneal injection of EAC cells after culturing in suspension (Sus) condition in presence of DMSO or PD98059. Black arrows depict macrometastatic nodules (a). Black circles indicate the presence of a micro metastatic lesion in histological sections (10× magnification) (b); n=3. G. Immunofluorescence of MDA-MB-231 cells cultured in suspension (Sus) condition for 24 hours in presence of vehicle control (DMSO) or AMPK-inhibitor (CC) and visualized with confocal microscopy (Z-stack, scale bar, 10 µM); n=3. Histograms represent the distribution of cells for pERK intensity (bin size 10). H. Box plots show distribution of expression of ERK, AMPK, or TFEB-dependent genes from RNA-sequencing data publicly available for 15 single CTCs pools and matched 14 CTC clusters isolated from ten breast cancer patients (SC, single CTCs; CL, CTC cluster) (<u>GSE51827</u>) (also see **Figure S12**). I. **Model:** Matrix-deprivation triggered AMPK inhibits ERK activity through phosphorylation of PEA15. Phosphorylation of PEA15 leads to inhibition of ERK activity, which results in increased TFEB level. Increased TFEB promotes autophagy maturation as well as AMPK activity. TFEB mediated upregulation of AMPK activity can enhance the inhibitory effect on ERK, resulting in increased autophagy maturation and, in turn, better cell survival in suspension.

Since AMPK negatively regulates ERK, and ERK inhibition improves autophagy maturation, therefore we next examined if ERK inhibition would counter the deleterious effects of AMPK inhibition on anoikis. As expected, Inhibition of AMPK abrogated the co-localization of LAMP2 with GFP-LC3 puncta (**Figures 6D & S11A**), whereas simultaneous ERK inhibition rescued this co-localization **(Figures 6D & S11A)**. We confirmed the same observation using electron microscopy (**Figure S11B**). Consistent with this observation, AMPK inhibition-mediated apoptosis was also partially rescued by co-treatment with PD98059 **(Figure 6C)**.

Taken together, these data highlight the role of ERK signaling downstream to AMPK activation in regulating autophagy maturation and anoikis.

Previously, we observed pAMPK^high^/pERK^low^ and pAMPK^low^/ pERK^high^ states of cells in matrix-deprived condition (**Figure 3**). We have also shown that cells with low ERK activity in matrix-deprived condition have better survival advantage than the cells with high ERK activity (**Figure 6E**). We further tested the significance of down modulation of ERK signaling in murine experimental metastasis model using Ehrlich Ascites Carcinoma (EAC) cells. Intraperitoneal injection of EAC cells that were treated with PD98059 led to increased number of metastatic nodules (**Figure 6F**), thus reinforcing our *in vitro* data. Additionally we assessed lung metastasis in BALB/c mice following tail-vein injection of eGFP expressing 4T1 cells injected, which also exhibit negative cross-talk between AMPK and ERK (**Figure S11C**). Consistent with our previous results, we observed a significantly higher number of lung nodules in mice injected with PD98059-treated cells (**Figure S11D**) which demonstrated that decreased ERK activity is better for survival in suspension and metastasis. Based on this, we hypothesized that pushing the cells from low ERK activity toward higher pERK state would promote anoikis. To achieve this, we broke AMPK-ERK feedback loop by AMPK inhibition and observed a shift in the cellular state from heterogeneous toward homogeneous high pERK population (**Figure 6G & S11E**). This increase in pERK population promotes anoikis due to blockage in autophagy maturation (**Figures 6C, 6D, & 6E**), which can be reverted by inhibition of ERK activity (**Figures 6C,6D, 6E & 6F**). Therefore, we suggest that inhibiting AMPK could help to reduce such non-genetic heterogeneity in the matrix-deprived cells and make these cells unfit to survive.

To further explore the biological relevance of the AMPK-ERK-TFEB axis in metastasis, we analysed the RNA-sequencing data of CTCs and CTC-clusters isolated from breast cancer patients (Aceto et al., 2014). The observed heterogeneity in ERK/AMPK activities and TFEB levels in matrix detached breast cancer cells in vitro was captured in vivo (**Figure S12**). Interestingly, we observed higher association of AMPK and TFEB gene signature with CTC-clusters (**Figures 6H, & S12A**) that were reported to corroborate with increased metastasis and poor survival (Aceto et al., 2014). Conversely, higher ERK gene signature was observed with single-CTCs (**Figures 6H & S12B**). These data are in corroboration with our in vitro studies where we have demonstrated that matrix-deprived breast cancer cells with pAMPK^high^/pERK^low^ status express higher levels of TFEB and show better survival.

## Discussion

Adaptation to matrix deprivation is fundamental for successful metastasis. Phenotypic heterogeneity and plasticity within different cell populations of cancer pose a major challenge to effective treatment strategy [32, 33]. The existence of such heterogeneity is beginning unveiled more frequently due to high-resolution techniques such as single-cell RNA-seq, and biosensor-based microscopic analysis [34–36], however, the origins and implications of such heterogeneity remain largely elusive. In this study, we demonstrate phenotypic (i.e. non-genetic) heterogeneity of ERK signaling in the matrix-deprived conditions. Here we have elucidated the mechanism for emergence of ERK heterogeneity – through feedback loop among AMPK, ERK and TFEB. We show that the AMPK^high^/ERK^low^/TFEB^high^ state enables overcoming autophagy maturation arrest, thereby facilitating anoikis resistance (**Figure 6I**). Thus, targeting this feedback loop might offer a novel and rational anti-cancer treatment strategy.

ERK signaling is often hyper-activated in cancers, leading to uncontrolled growth [37]. However, accumulating evidence also suggests that sustained activation of ERK may promote apoptosis [38]. Even though we see an increase in ERK activity in a population of matrix-deprived cells at early time point (24 hr), but colonies formed by long-term (one week) culture of cells in suspension show decreased ERK activity. This was an unexpected finding as ERK is often associated with anoikis resistance [39, 40]. Upon further analysis at single cell level, we observed heterogeneity in the population for ERK activity. Isolation of ERK^high^ and ERK^low^ population based on promoter-reporter system revealed that cells in suspension with low ERK activity survive better than the counter population. Recent reports also suggest that cells with low ERK activity are more sturdy and tolerate drug therapy [41], show more stem cell-like properties and survive better to form anchorage-independent colonies [42]. Similarly, we lately demonstrated that inhibition of Akt, typically a pro-tumorigenic signaling molecule [43], confers better survival benefits to matrix-detached cells [9]. Altogether, these data begin to unfurl novel contextual signaling networks that can maintain a pro-survival state of matrix-detached cancer cells.

In cancer, autophagy plays dual roles as cancer suppressor (during cancer initiation) as well as in tumor progression (especially in the advanced stage of cancer) [44]. Initially, autophagy was thought to be a tumor-suppressive mechanism, based on observations that *Beclin1* was deleted in most breast cancers and its overexpression in MCF7 cells reduced the tumorigenesis [45, 46]. Similarly, deletion of *Atg5* and *Atg7* in mice led to development of benign tumor [47, 48]. However, autophagy is robustly activated in tumor cells facing stresses such as oncogenic insult, starvation, hypoxia, matrix deprivation, or higher metabolic demands [48]. We show that cells with high ERK activity in suspension display autophagy maturation block along with autophagy induction. Induction of autophagy but maturation blockage is more challenging for cell survival compared to only blockage of the autophagic maturation [15]. We also observed high cleaved-caspase levels in the cells with autophagy blockage and high ERK activity, where ERK activation is responsible for inhibition of autophagy maturation. In conclusion, we find that cells in matrix-deprived condition have heterogeneity in autophagy maturation which can be regulated by ERK activity.

AMPK can activate as well as inhibit ERK activity in different contexts [49, 50]. Our finding uncovers AMPK as an inhibitor of ERK activity and maintains heterogeneous population of ERK^high^/AMPK^low^ and vice versa. We found similar inverse corelation between AMPK and ERK signaling in the detached cells from lumen the lactating mammary gland, patient-derived breast cancer tissue as well as in CTCs. We previously reported that matrix-deprivation triggered AMPK phosphorylates PEA15 [8] - a scaffold protein for ERK [51]. The phosphorylation of PEA15 targets ERK to one of its substrates, RSK2, thus promoting its activity [52]. We show here that the heterogeneity in ERK activity is maintained by AMPK-mediated phosphorylation of PEA15 at S116 position. AMPK, by phosphorylating PEA15 at S116, inhibits the interaction of MEK, upstream kinase of ERK, with PEA15-ERK complex, thereby hindering ERK activity. However, we did not observe change in MEK activity under matrix-deprived condition. Previous literature using yeast two-hybrid system showed that among other members of the MAPK signaling pathway, only ERK interacts with PEA15 [53]. Our immunoprecipitation studies revealed the presence of MEK in PEA15-pull down complex in matrix-deprived cells. There could be a direct interaction between MEK and PEA15, aided possibly by post-translational modifications under matrix detachment. Alternatively, this interaction could be indirect via ERK which interacts directly with PEA15 irrespective of the phosphorylation status [29]. Our data show that AMPK-mediated phosphorylation of PEA15 contributes to cancer cell survival in suspension through inhibiting ERK activity and promoting autophagy maturation.

ERK regulates autophagy by restricting the nuclear entry of TFEB - a master transcription factor for genes involved in lysosomal biogenesis and autophagy [30]. We observed an increase in TFEB expression in suspension which was heterogeneous and inversely correlated with ERK activity. We did not see change in TFEB nuclear localization upon ERK inhibition in the matrix-deprived condition. Another MiTF-TFE family transcription factor, TFE3 also does not show dependency on ERK signaling in the matrix-deprived condition, suggesting increase in TFEB level is dependent and specifically regulated by ERK mediated signaling. ERK inhibition did not change the transcript levels of TFEB, but led to its increased stability, suggesting a post-transcriptional control of TFEB by ERK. Interestingly, increase in TFEB level needs activation of AMPK, which is upstream of ERK. The exact mechanism for the increase in TFEB level under matrix-deprived condition is unclear and will be investigated in the follow-up study. However, mass-spec data suggests that the stability of TFEB might be regulated by altering SUMO3 under matrix deprived condition.

We find a positive feedback loop between AMPK and TFEB involving ERK. This feedback loop can generate heterogeneous subpopulation of cells with pAMPK^high^/pERK^low^/ TFEB^high^ or pAMPK^low^/pERK^high^/TFEB^low^ status, also suggested by mathematical modelling. The pAMPK^high^/pERK^low^/ TFEB^high^ cells displayed elevated autophagic maturation, less apoptosis, and increased sphere-forming potential compared to pAMPK^low^/pERK^high^/TFEB^low^ cells. Supporting this, we observed more metastasis in mice injected with EACs cells intraperitoneally after ERK inhibition. This has been reported also in a recent report that showed, mammary tumor cells with pERK^high^ status were less tumorigenic when cultured in matrix-deprived condition, as compared to pERK^low^ cells [42]. Importantly, we can push pAMPK^high^/pERK^low^/ TFEB^high^ (More fit) to pAMPK^low^/pERK^high^/TFEB^low^ (less fit) by targeting AMPK in the signalling node to prevent metastasis. Finally, in publicly available RNA-seq data of breast cancer patients [54], we observed an elevated AMPK and TFEB signature but lower ERK signature activity in CTCs clusters relative to that in individual CTCs. This observation corroborates with higher aggressiveness and poor prognosis of clusters of CTCs [54], and emphasizes a pro-metastatic role of pAMPK^high^/pERK^low^/TFEB^high^ state. Collectively, our data suggest that pAMPK^high^/ pERK^low^/TFEB^high^ status is favorable for survival under matrix deprivation.

The exitance of heterogeneous population and their significance in cancer cell survival in the matrix-deprived conditions are largely unknown. In our study, we uncovered the existance of non-genetic and plastic heterogeneity in the matrix-deprived cancer cells. Cancer cells with pAMPK^high^/pERK^low^/TFEB^high^ show better survival with respect to its counter population. AMPK, ERK, TFEB can cross talk in a complex but regulated manner. Mutually activating – such as those between phosphorylated AMPK and TFEB – or mutually inhibiting – such as those operating between phosphorylated AMPK and phosphorylated Akt [9] – feedback loops can facilitate phenotypic plasticity among these subpopulations [55], thus generating non-genetic heterogeneity [56] due to underlying multistability [57]. Similar feedback loops have been reported to mediate cellular plasticity in cancer cells such as epithelial-mesenchymal plasticity [58, 59], switching between a cancer stem cell and a non-cancer stem cell [60], metabolic plasticity [57, 61], or switching between a matrix-deprived and a matrix-attached condition [9]. Such dynamic interconversions – driven by multiple forms of ‘noise’ or stochasticity in biological systems [62, 63] may enable a more adaptive cellular stress response. Some feedback loops may also operate across multiple cells, hence affecting these different processes in a non-cell autonomous manner, and giving rise to intriguing spatial patterns of heterogeneity [64]. Therefore, identifying and breaking these feedback loops may severely impair non-genetic heterogeneity and consequently the fitness of a stressed cellular population.

## Acknowledgements

We thank Prof. Wilfried Roth for kindly providing the plasmids pcDNA3-Flag-WT-PEA15 and S116A-PEA15. We would like to acknowledge Dr. Benoit Viollet for AMPK DKO cells. This work was majorly supported from Wellcome Trust/DBT India Alliance Fellowship (grant number 500112-Z-09-Z) awarded to A. Rangarajan. MKJ acknowledges Ramanujan Fellowship provided by SERB, DST, Government of India (award number SB/S2/RJN-049/2018). We acknowledge support from DBT-IISc partnership program to AR, and DST-FIST and UGC, Government of India, to the Department of MRDG. SK acknowledges Council for Scientific and Industrial Research for CSIR fellowship (18-12/2011 (ii) EU-V). We would like to thank FACS-facilities (IISc and MRDG), and Central animal facility (IISc).

