## Supplemental data for "Feedback loop involving AMPK, ERK and TFEB generates non-genetic heterogeneity in matrix-deprived cancer cells and regulates survival"

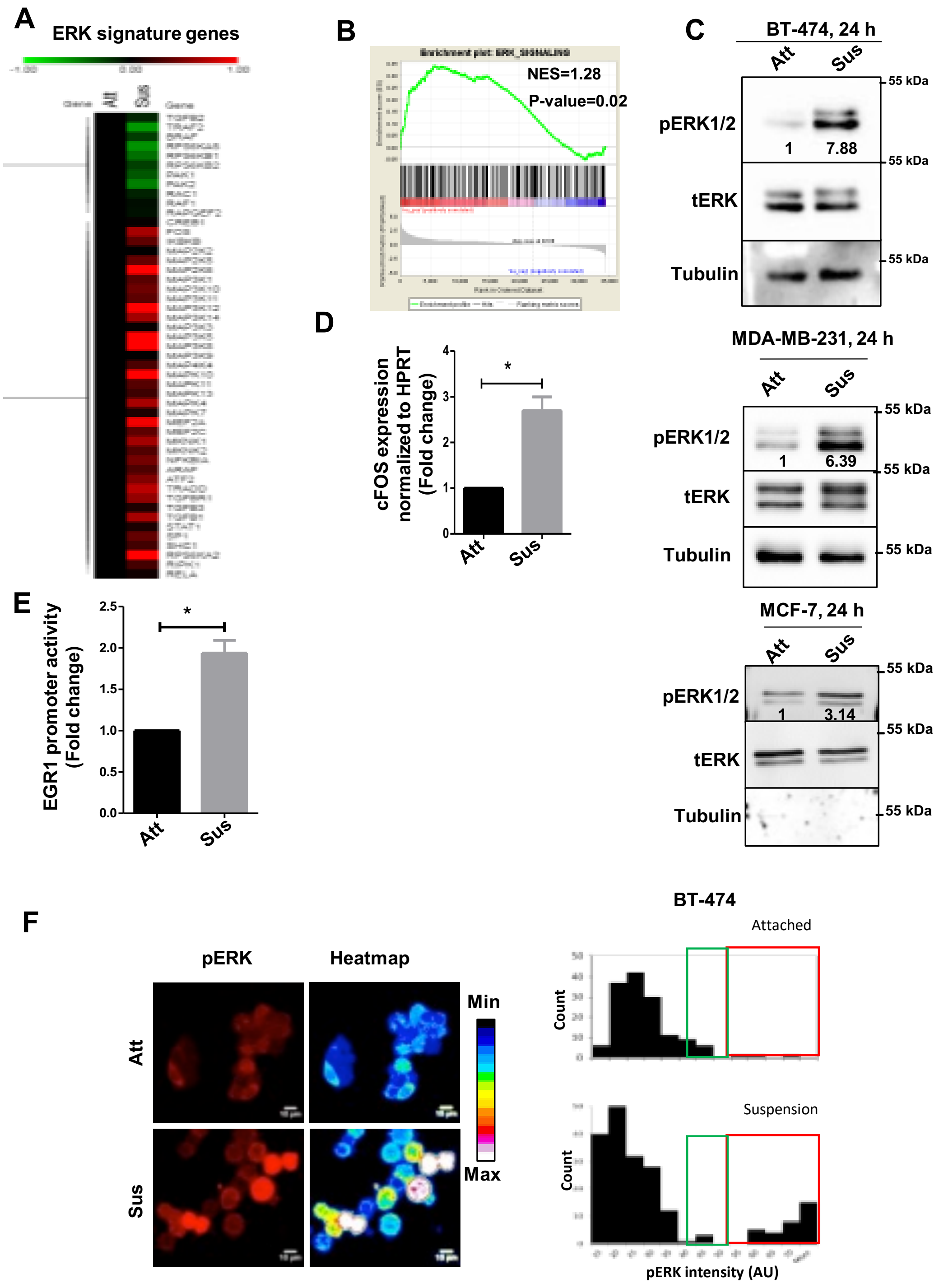


**Figure S1. Increase in ERK signaling and heterogeneity in matrix-deprived conditions**

A & B). Comparative analysis of gene expression from microarray data of MDA-MB-231 cells cultured in suspension (Sus) versus attached (Att) conditions for 24 hours:

(A). Heat map of semi-supervised clustering of ERK pathway signature genes. Red represents highly expressed genes, while green represents downregulated genes.

(B). Gene Set Enrichment Analysis (GSEA) plot for ERK pathway.

C. Representative immunoblots of BT-474, MDA-MB-231, and MCF-7 cells cultured in attached (Att) or suspension (Sus) conditions for 24 hours; n=3.

D. MDA-MB-231 cells were cultured in attached (Att) or suspension (Sus) conditions for 24 hours and harvested for qRT-PCR; n=3.

E. Graph represents MDA-MB-231 cells stably expressing EGR1(promoter)-TurboRFP cultured in attached (Att) and suspension (Sus) conditions for 24 hours and harvested for analysis of RFP intensity by flow cytometry; n=3.

F. Fluorescent images of BT-474 cells cultured in attached (Att) or suspension (Sus) conditions for 24 hours, probed for pERK, and visualized by confocal microscopy (Z-stack, scale bar, 20μM); n=3. Heat map was generated with the help of ImageJ. Histograms represent the distribution of cells cultured in attached (Att) or suspension (Sus) conditions for 24 hours based on pERK intensity.

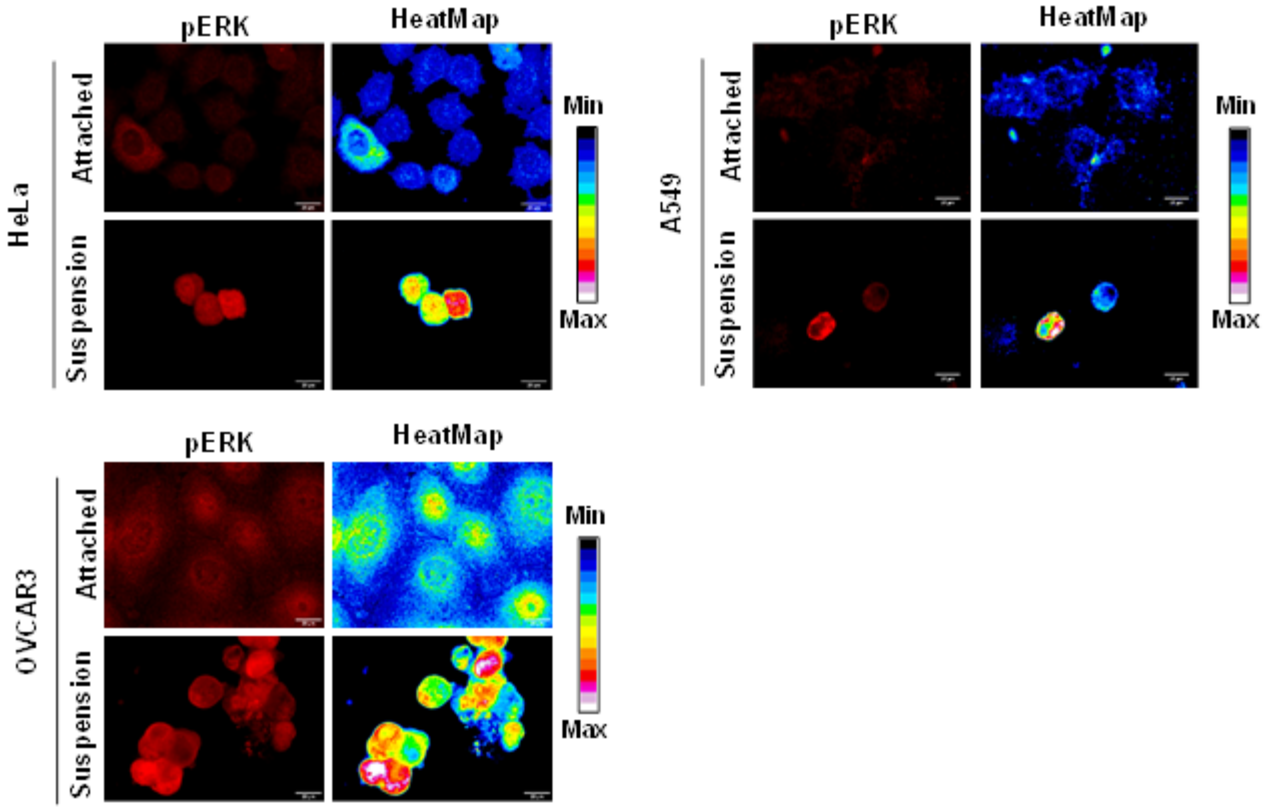


**Figure S2. Increased ERK signaling in matrix-deprived condition**

Fluorescent images of HeLa, A549, OVCAR3 cells cultured in attached (Att) or suspension (Sus) conditions for 24 hours, probed for pERK, and visualized by confocal microscopy (Z-stack, scale bar, 20μM); n=3. Heat map was generated with the help of ImageJ.

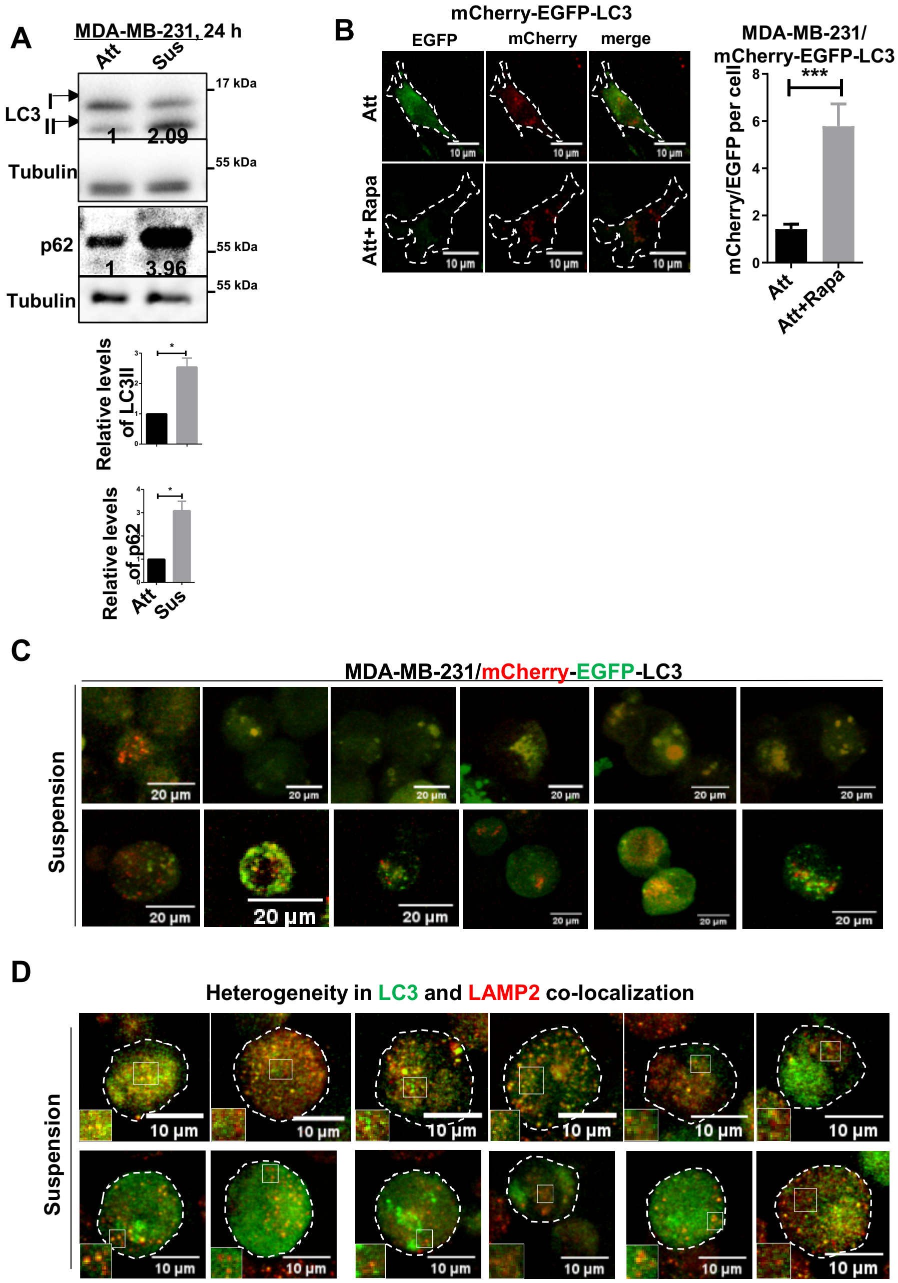


**Figure S3. Autophagy heterogeneity in matrix-deprived conditions**

A. Immunoblot analysis of MDA-MB-231 cells cultured in attached (Att) or suspension (Sus) conditions for 24 hours. Graphs represent densitometric quantification of immunoblots; error bars, mean±SEM; n=3.

B. Fluorescent images of MDA-MB-231 cells stably expressing mCherry-EGFP-LC3, cultured in attached (Att) conditions with or without rapamycin (Rapa) for 24 hours and visualized with confocal microscopy (Z-stack, scale bar, 20μM); n=3. Graph represents ratio of mChrrey/EGFP intensity.

C. Fluorescent images of MDA-MB-231 cells stably expressing mCherry-EGFP-LC3, cultured in suspension condition for 24 hours and visualized with confocal microscopy (Z-stack, scale bar, 20μM); n=3.

D. Fluorescent images of MDA-MB-231 cells cultured in attached (Att) or suspension (Sus) conditions for 24 hours, probed for LC3 and LAMP2, and visualized by confocal microscopy (Z-stack, scale bar, 10 μM); n=2.

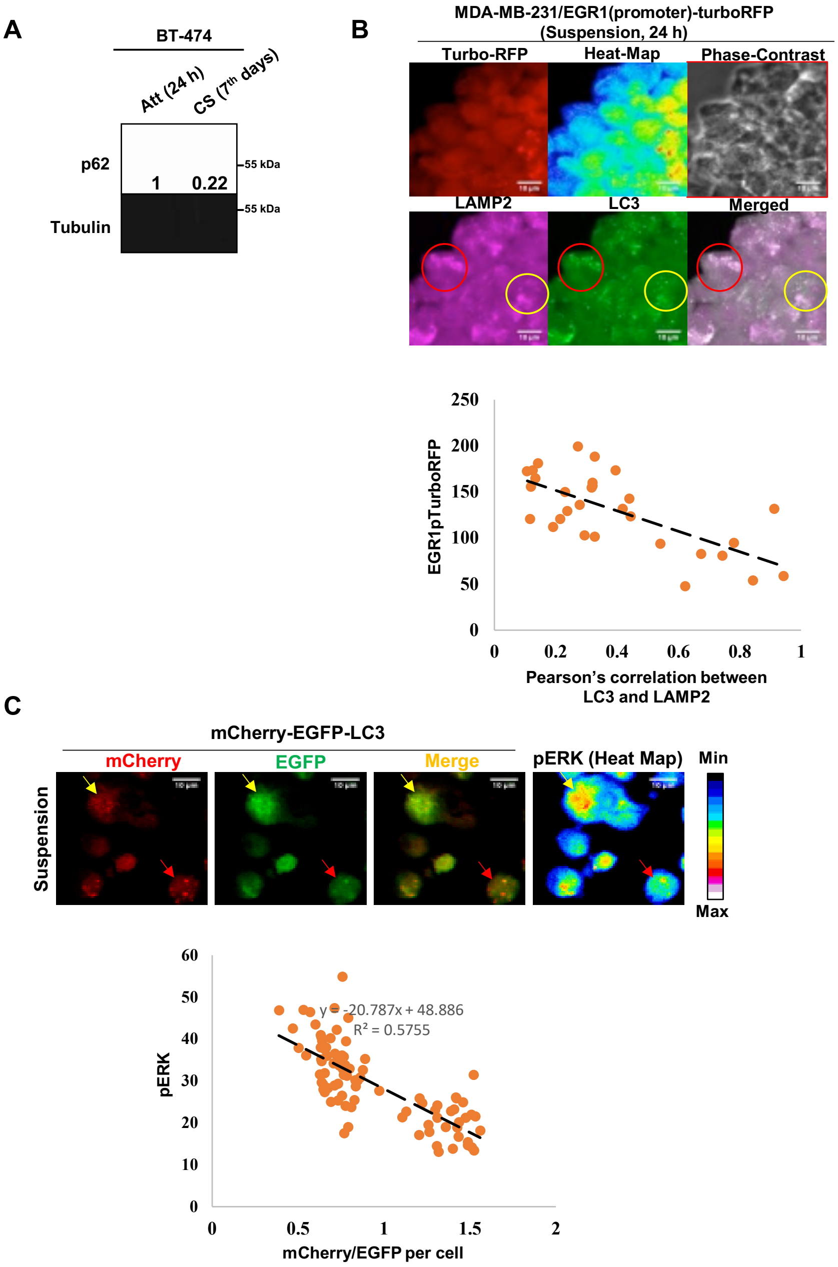


**Figure S4. Inverse corelation between ERK status and autophagy maturation in matrix-deprived conditions**

A. Representative immunoblots of BT-474 cells cultured in attached (Att, 24 hours) or anchorage independent cancer sphere (CS, 7 days) in the absence of serum; n=3.

B. Immunofluorescence of MDA-MB-231 cells stably expressing $EGR1_{(promoter)}$-TurboRFP with anti-LAMP2 antibody (Cy3) and co-localization with LC3 puncta (Alexa fluor-488) in cells cultured in suspension (Sus) condition for 24 hours and visualized with confocal microscopy (Z-stack, scale bar, 10 μM); n=2. Red circle represents $ERK^{low}$ and co-localization of LAMP2 and LC3, yellow circle represents $ERK^{high}$ and less co-localization of LAMP2 and LC3. Graph represents co-relation between RFP intensity and Pearson's correlation between LC3 and LAMP2; colocalization of LAMP2 and LC3 was measured by Pearson's correlation coefficient employing Coloc-2 plugin in ImageJ and represented as dot plot (each dot represents single cell).

C. Fluorescent images of MDA-MB-231 cells cultured suspension (Sus) condition for 24 hours, probed for pERK, and visualized by confocal microscopy (Z-stack, scale bar, 10 μM); n=3. Heat map was generated with the help of ImageJ. Graph represents co-relation between pERK intensity per cell with its own ratio of mChrrey/EGFP intensity (each dot represents single cell).

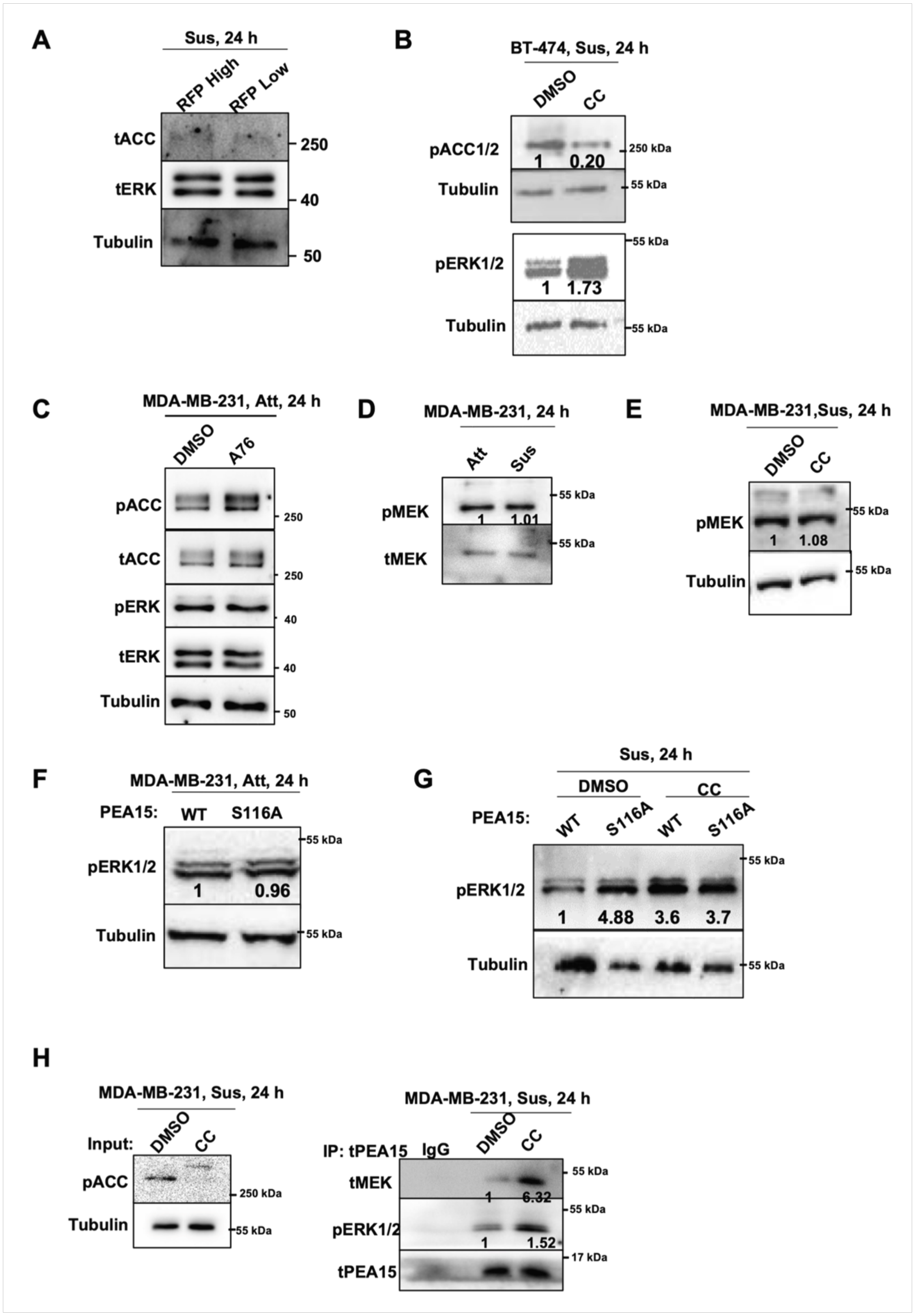


**Figure S5. AMPK inhibits ERK through PEA15-phosphorylation**

A-F). Immunoblots of following cell lysates were harvested and probed for specified proteins:

(A). MDA-MB-231 cells stably expressing $EGR1_{(promoter)}$-TurboRFP were cultured in suspension (Sus) condition for 24 hours. High and low RFP subpopulations were separated by FACS sorting; n=3.

(B). BT-474 cells cultured in suspension (Sus) condition for 24 hours in the presence of vehicle control (DMSO) or AMPK inhibitor (CC); n=3.

(C). MDA-MB-231 cells cultured in attached condition for 24 hours and treated with vehicle control (DMSO) or AMPK inhibitor (CC) for indicated time; n=3.

(D). MDA-MB-231 cells cultured in attached (Att) or suspension (Sus) conditions for 24 hours; n=3.

(E). MDA-MB-231 cells cultured in suspension (Sus) condition for 24 hours in the presence of vehicle control (DMSO) or AMPK inhibitor (CC); n=3.

(F). MDA-MB-231 cells stably overexpressing Flag-tag WT-PEA15 or S116A-PEA15 cultured in attached (Att) condition for 24 hours; n=3.

(G). MDA-MB-231 cells stably overexpressing Flag-tag WT-PEA15 or S116A-PEA15 cultured in suspension (Sus) condition for 24 hours in presence of vehicle control (DMSO) or AMPK inhibitor (CC); n=3.

H. Immunoblot analysis of immunoprecipitated (IP) product with IgG control or anti-Flag antibodies from MDA-MB-231 cells stably overexpressing Flag-tag WT-PEA15 cultured in suspension condition for 24 hours in presence of vehicle control (DMSO) or AMPK inhibitor (CC). 2% of the whole-cell lysate was used as input and probed for specified proteins; n=3.

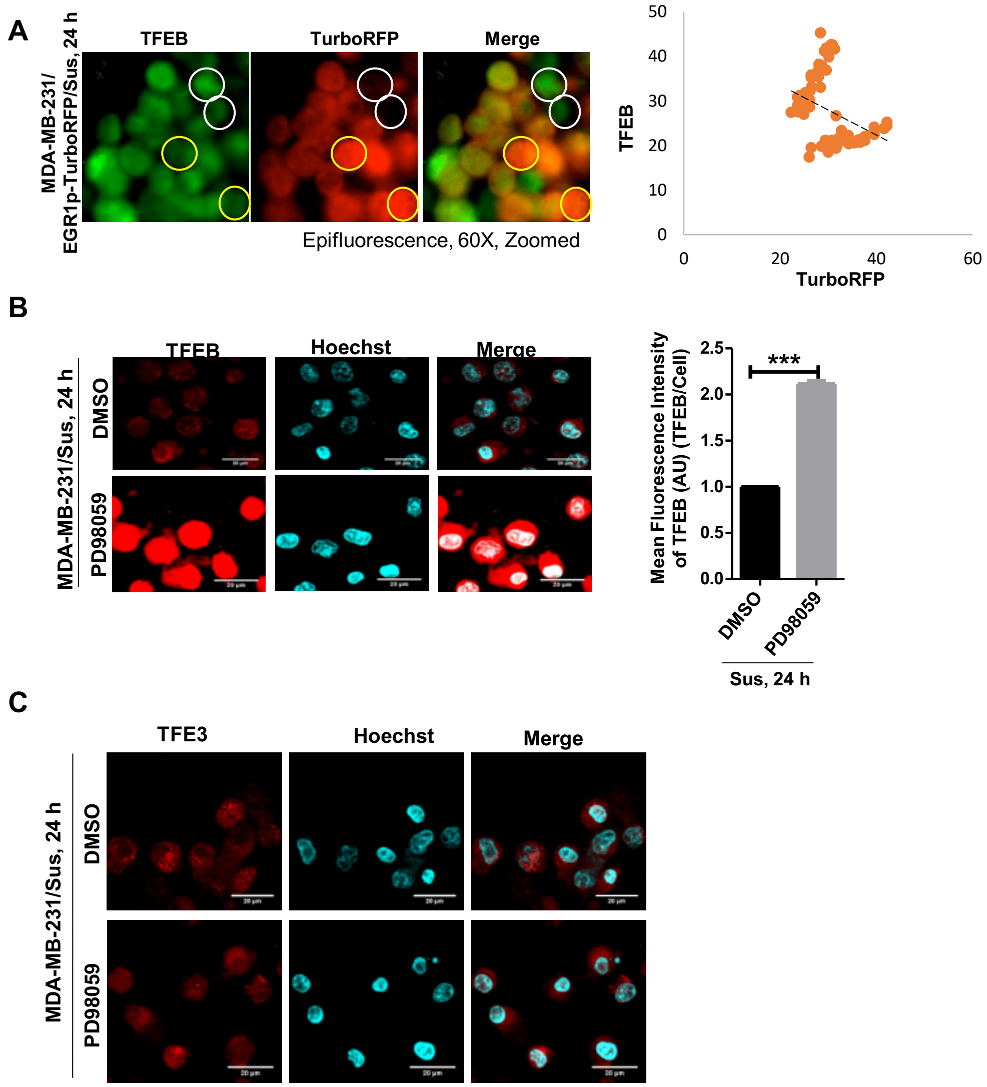


**Figure S6. Inverse co-relation between ERK activity and TFEB level**

A. Immunofluorescence of MDA-MB-231 cells stably expressing EGR1$_{(promoter)}$-TurboRFP with anti-TFEB antibody in cells cultured in suspension (Sus) condition for 24 hours and visualized with confocal microscopy (Z-stack, scale bar, 10 μM); n=3. Yellow circle area represent cell with TFEB$^{low}$/ERK$^{high}$ and white circle represents cells with TFEB$^{high}$/ERK$^{low}$ status. Co-relation graph was plotted using excel.

B. Immunofluorescence of MDA-MB-231 cells cultured in suspension (Sus) condition for 24 hours in presence of vehicle control (DMSO) or MEK-inhibitor (PD98059), probed for TFEB, and visualized by confocal microscopy (Z-stack, scale bar, 20 μM); n=3.

C. Immunofluorescence of MDA-MB-231 cells cultured in suspension (Sus) condition for 24 hours in presence of vehicle control (DMSO) or MEK-inhibitor (PD98059), probed for TFE3, and visualized by confocal microscopy (Z-stack, scale bar, 20 μM); n=2.

A

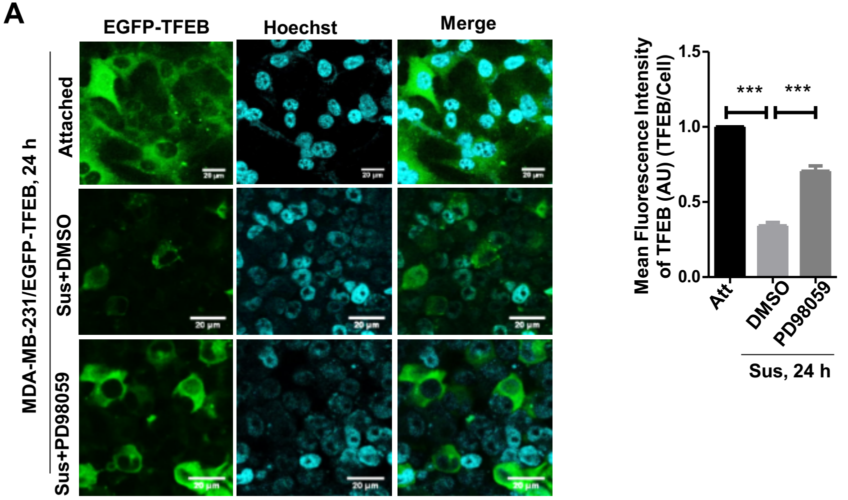


B

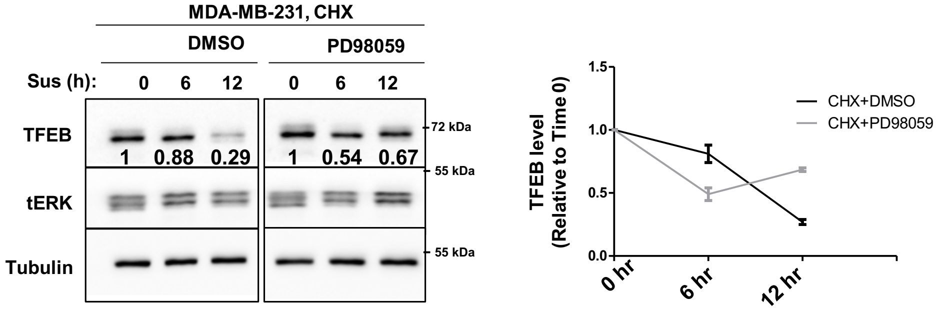


C

| Protein FDR confidence | Accession | Description | GFP-TFEB_Sus+DMSO/ GFP_Sus | GFP-TFEB_Sus +CC/GFP_Sus | GFP-TFEB_Sus+ CC/GFP-TFEB_Sus+ DMSO |
|---|---|---|---|---|---|
| High | P31947 | 14-3-3 protein sigma OS=Homo sapiens OX=9606 GN=SFN PE=1 SV=1 | 3.34 | 11.62 | 3.457627 |
| High | A8MU27 | Small ubiquitin-related modifier 3 OS=Homo sapiens OX=9606 GN=SUMO3 PE=1 SV=1 | 1.505 | 3.741 | 2.491935 |
| High | P05161 | Ubiquitin-like protein ISG15 OS=Homo sapiens OX=9606 GN=ISG15 PE=1 SV=5 | 0.054 | 0.136 | 2.461538 |
| High | P22314 | Ubiquitin-like modifier-activating enzyme 1 OS=Homo sapiens OX=9606 GN=UBA1 PE=1 SV=3 | 0.796 | 1.849 | 2.323671 |
| High | O14933 | Ubiquitin/ISG15-conjugating enzyme E2 L6 OS=Homo sapiens OX=9606 GN=UBE2L6 PE=1 SV=4 | 0.086 | 0.194 | 2.238095 |
| High | P62979 | Ubiquitin-40S ribosomal protein S27a OS=Homo sapiens GN=RPS27A PE=1 SV=2 | 1.719 | 3.213 | 1.870283 |

**Figure S7. ERK negatively regulates TFEB level**

A. Fluorescence images of MCF-7 cells transiently expressing EGFP-TFEB cultured in suspension (Sus) condition for 24 hours in presence of vehicle control (DMSO) or MEK-inhibitor (PD98059); n=3.

B. Immunoblots analysis of MDA-MB-231 cells pre-treated with vehicle control (DMSO) or MEK-inhibitor (PD98059) were treated with cycloheximide (CHX) for 30 minutes, followed by suspension (Sus) condition for indicated time points. Graph represents quantification of TFEB normalized to Tubulin level; n=2.

C. List of altered ubiquitin related proteins detected using quantitative Mass-spectrometry of MCF7 cells stably expressing EFGF-TFEB cultured in suspension (Sus) condition for 24 hours in presence of vehicle control (DMSO) or MEK-inhibitor (PD98059). n=2.

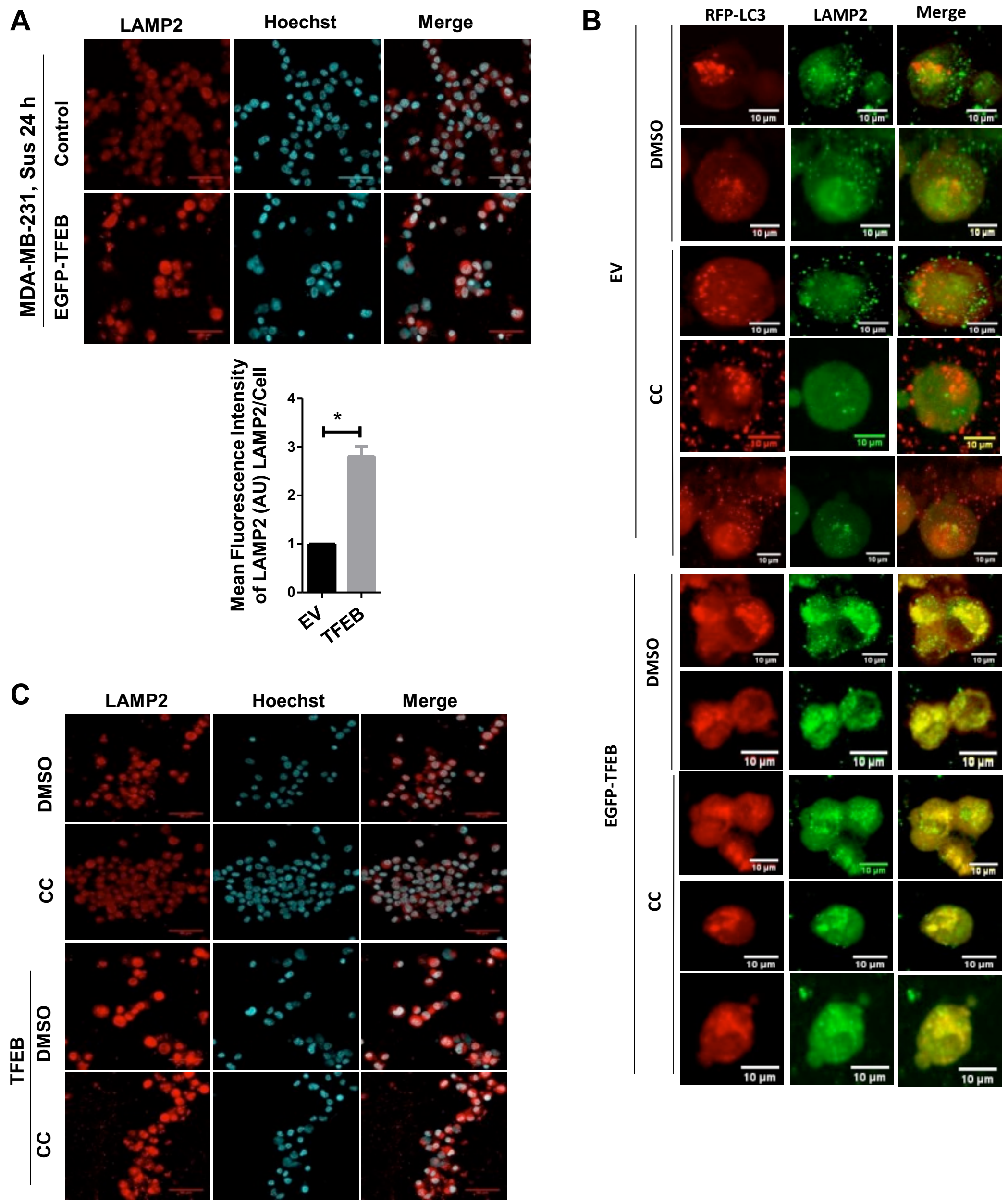


**Figure S8. TFEB can rescue phenotypes due to AMPK-inhibition**

Immunofluorescence images of (A). MDA-MB-231 cells stably overexpressing control empty vector or EGFP-TFEB were cultured in suspension (Sus) condition for 24 hours with anti-LAMP2 antibody (Cy3) and visualized with confocal microscopy (≥50 cells analysed per sample/experiment); n=3. (B). MDA-MB-231 cells stably overexpressing control empty vector or EGFP-TFEB were cultured in suspension (Sus) condition for 24 hours in presence of vehicle control (DMSO) or AMPK inhibitor (CC) with anti-LAMP2 antibody (Alexa 488) and co-localization with RFP-LC3 puncta. Representative fluorescent images are visualized with confocal microscopy (Z-stack, scale bar, 10 μM); n=3 (C). MDA-MB-231 cells stably overexpressing control empty vector or EGFP-TFEB were cultured in suspension (Sus) condition for 24 hours in presence of vehicle control (DMSO) or AMPK inhibitor (CC) with anti-LAMP2 antibody (Alexa 488). Representative fluorescent images are visualized with confocal microscopy (Z-stack, scale bar, 10 μM); n=3

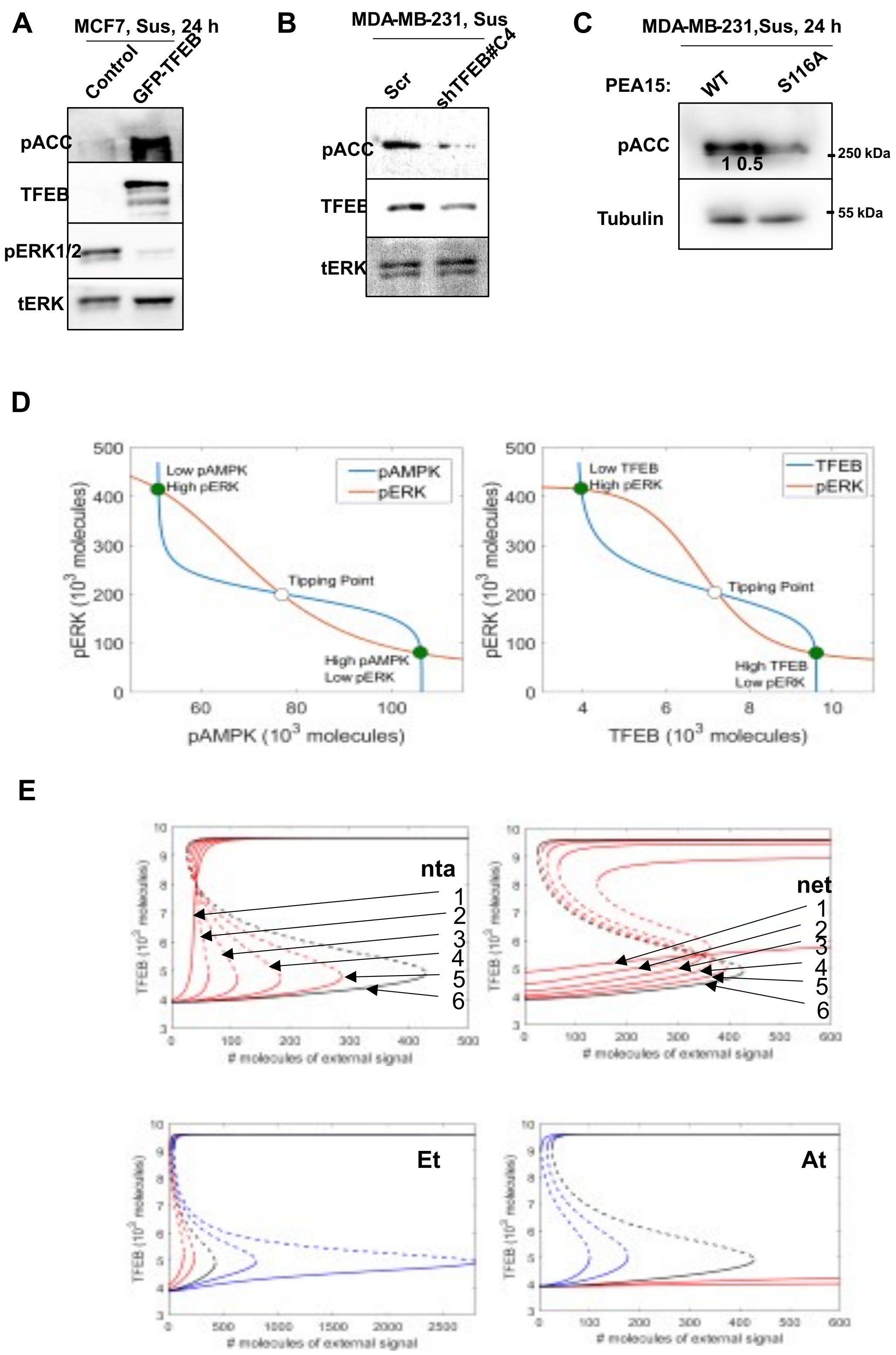


**Figure S9. Cellular plasticity in the matrix-deprived conditions**

A-C). Immunoblots analysis of (A). MDA-MB-231 cells stably overexpressing control empty vector or EGFP-TFEB were cultured in suspension (Sus) condition for 24 hours; (B). MDA-MB-231 cells stably overexpressing Scr control or shTFEB#C4 cultured in suspension (Sus) condition for 24 hours; (C). MDA-MB-231 cells stably overexpressing Flag-tag WT-PEA15 or S116A-PEA15 cultured in suspension (Sus) condition for 24 hours. n=3.

D. Nullclines for pERK-pAMPK and pERK-TFEB axes, similar to that in Fig 8C, with stable steady states (green circles) and tipping points (white circles)

E. Sensitivity analysis for model parameters (Hill coefficients and total protein concentrations of ERK and AMPK – Et and At). In all the plots, black curves represent the system with the original parameter set, blue curves represent an increased value of the parameter of interest, and red curves represent a decreased value of the parameter. The Hill coefficients were varied from 1 to 6 (labelled in the Figure) and At and Et were varied as 80%, 90% (red) and 110%, 120% (blue) of the actual values considered in the model. Dotted curves shown unstable states and solid curves show stable steady states.

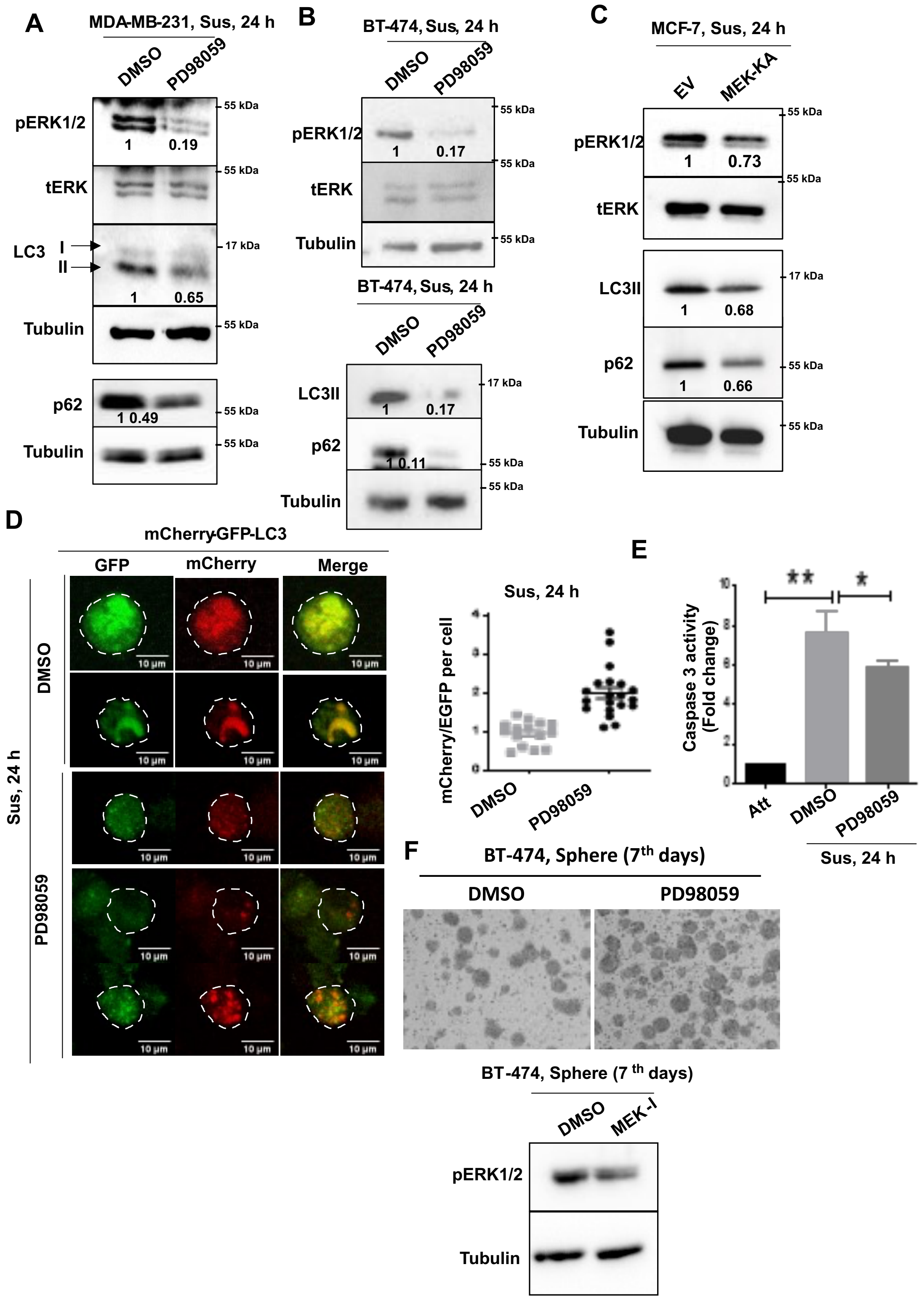


**Figure S10: ERK negatively regulates autophagy maturation in matrix deprived conditions**

A-C). Immunoblots analysis of (A). MDA-MB-231, (B). BT-474 and (C). MCF7 cells cultured in suspension (Sus) condition for 24 hours in presence of vehicle control (DMSO) or MEK-inhibitor (PD98059) for 24 hours; n=3.
D. Fluorescence images of MDA-MB-231 cells stably expressing mCherry-EGFP-LC3 cultured in suspension (Sus) condition for 24 hours in presence of vehicle control (DMSO) or MEK-inhibitor (PD98059); Graph represents ratio of mChrrey/EGFP intensity; n=3.

E. MDA-MB-231 cells cultured in attached (Att) and suspension (Sus) condition for 24 hours in presence of vehicle control (DMSO) or MEK-inhibitor (PD98059) and harvested for flow cytometric analysis of caspase-3 activity; n=3.

F. BT-474 cells cultured in suspension condition (Sus) or anchorage independent cancer spheres (CS) in methylcellulose for 7-days; imaged and harvested for immunoblotting.

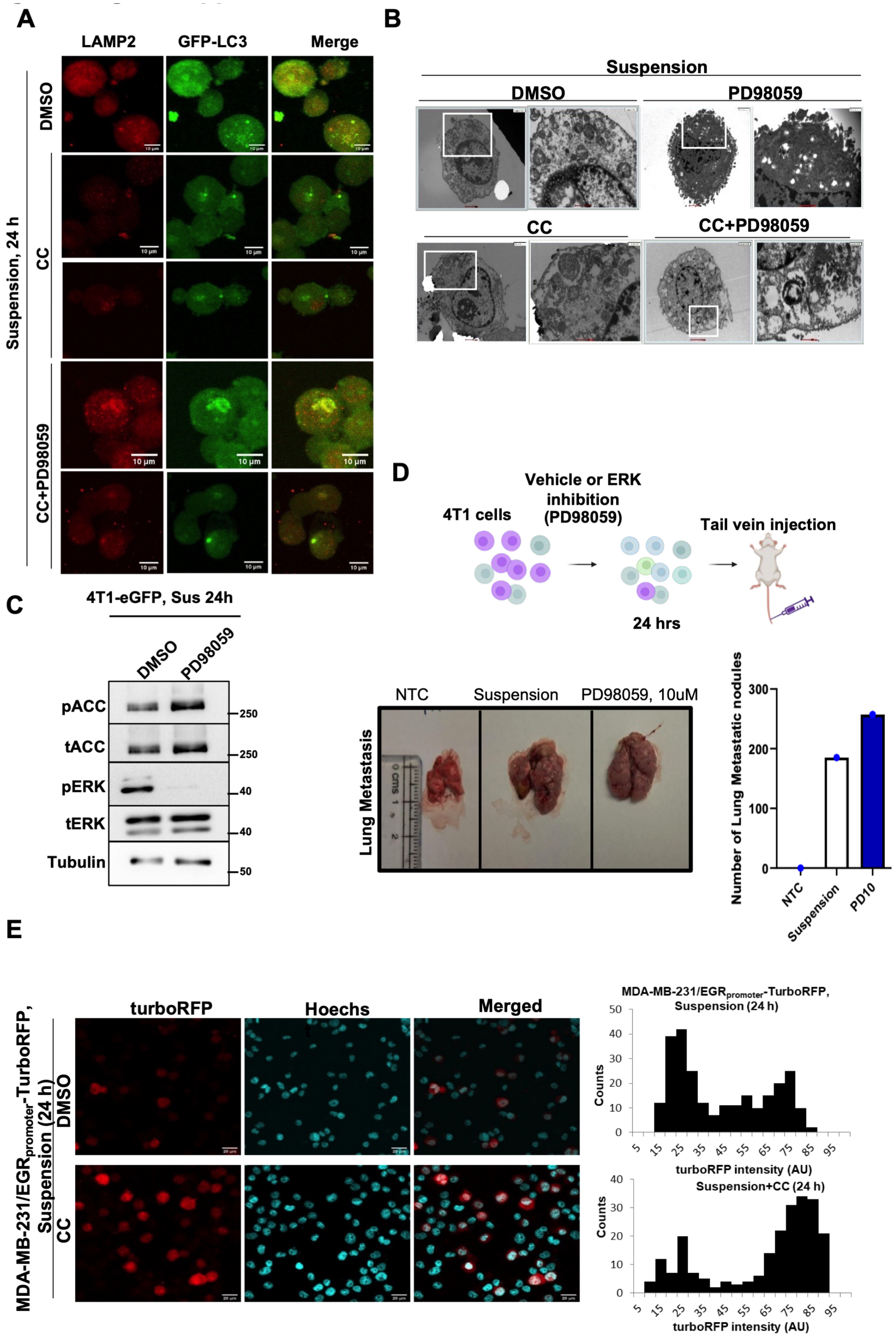


**Figure S11: AMPK mediated inhibition of ERK activity promotes autophagy maturation**

A & B). MDA-MB-231 cells cultured in suspension (Sus) condition for 24 hours in presence of vehicle control (DMSO) or AMPK inhibitor (compound C, CC) with or without MEK-inhibitor (PD98059) and harvested for (A). immunofluorescence and (B). electron micrographs analysis; n=3.

C. 4T1-eGFP cells cultured in suspension (Sus) condition for 24 hours in presence of vehicle control (DMSO) or MEK-inhibitor (PD98059) for 24 hours and harvested for immunoblotting; n=3

D. Representative schematic of experimental plan, images and quantification of lung metastastatic nodules following tail vein injection of 4T1-eGFP cells after culturing in suspension (Sus) condition in presence of DMSO or PD98059; n=3.

E. Fluorescence images of MDA-MB-231 cells stably expressing $EGR1_{(promoter)}$-TurboRFP cultured in suspension (Sus) condition for 24 hours in presence of vehicle control (DMSO) or AMPK inhibitor (compound C); Histograms represent the distribution of cells for RFP intensity (bin size 10).; n=3.

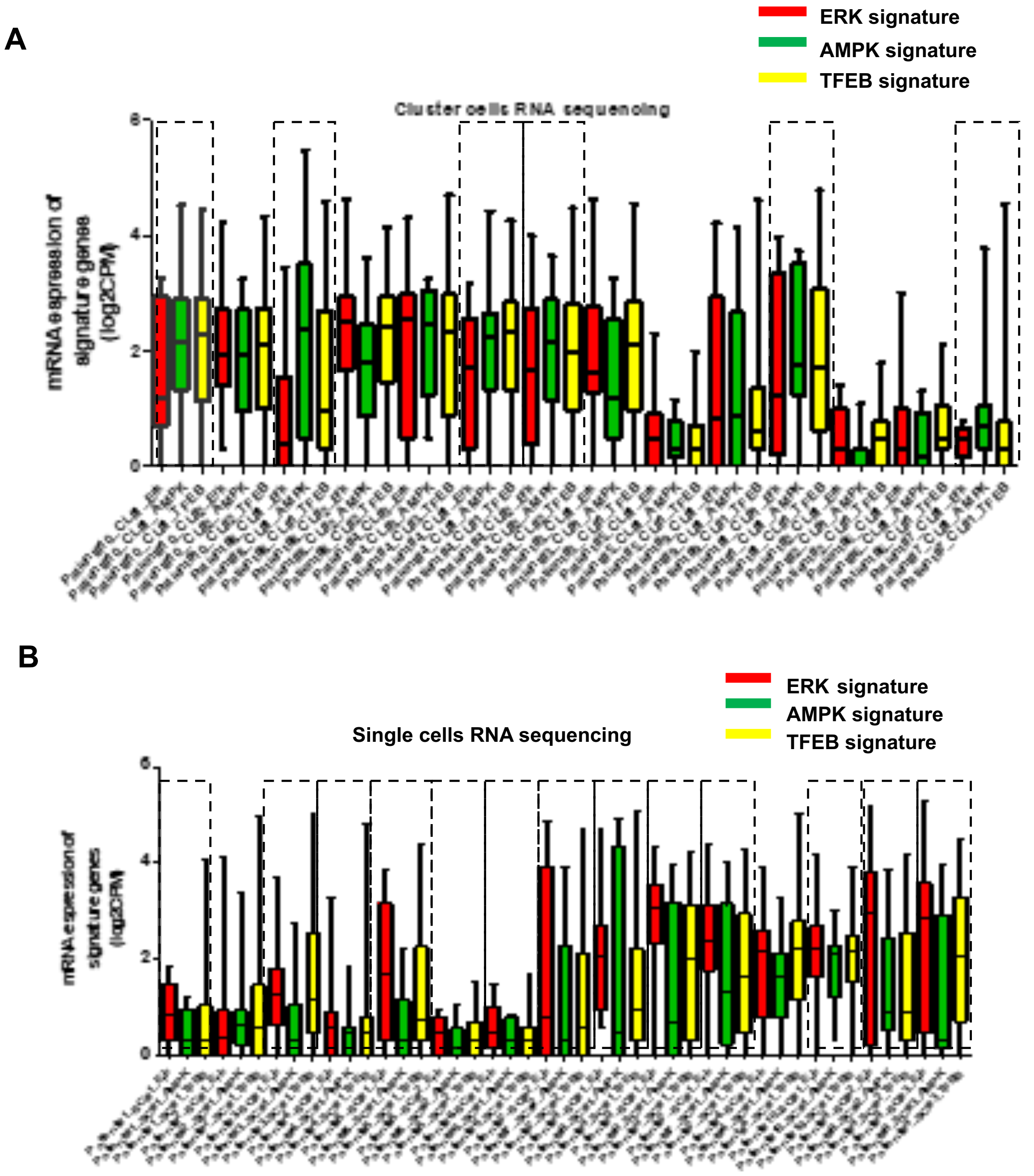


**Figure S12: RNA-seq data analysis from single and clustered CTCs**

A & B). Box plots show individual distribution of gene expression of ERK, AMPK, or TFEB-dependent genes from RNA-sequencing data publicly available in 14 CTC clusters (A) and matched 15 single CTCs (B) pools isolated from ten breast cancer patients (SC, single CTCs; CL, CTC cluster)
