## Supplementary material for "Feedback loop involving AMPK, ERK and TFEB generates non-genetic heterogeneity in matrix-deprived cancer cells and regulates survival": Method and materials

**Cell culture and transfection**

Breast cancer cell lines MDA-MB-231, MCF7, BT-474 and 4T1 (procured from ATCC in 2016 and validated by STR analysis) Cells were trypsinized and cultured for indicated time points on tissue culture dishes coated with 1% noble agar (Sigma-Aldrich) to mimic ECM-detachment Pharmacological agents were added immediately after trypsinization and prior to plating onto noble agar coated dishes. Long-term anchorage independent (AI) colony formation assay was performed by mixing 1x10^5^ cells with 1.5% methylcellulose and layered on top of 1.5 % noble agar coated dishes.

Cell transfection was performed using Lipofectamine 2000 according to manufacturer’s protocol. OptiMEM media was used for the transfection and changed to regular media 6 hours post transfection. Drugs (Puromycin or G418) for selection were used for generation of stable cells after 24 hours of transfection. MDA-MB-231 cells stably expressing mCherry-EGFP-LC3, GFP-LC3, EGR_(promoter)_-TurboRFP, pEGFP-TFEB, Scr (Control), or shTFEB#C5 were generated by transfection followed by FACS sorting.

**Pharmacological compounds**

Pharmacological compounds used in the study include compound C (10mM; Calbiochem), PD98059 (10 µM; CST), and rapamycin (100nM; Sigma-Aldrich). Dimethyl sulfoxide (DMSO, Thermo Scientific) was used as vehicle control for all the compounds except rapamycin, which was dissolved in ethanol.

**Caspase 3-activity assay**

Briefly, 1 x 10^6^ cells were incubated with 1 µL of Red-DEVD-FMK for 30 minutes at 37°C incubator maintaining 5% CO_2_. Caspase-3 activity was analysed by BD FACS-CantoII (Becton & Dickinson) containing a 488-nm Coherent Sapphire Solid State laser. Red fluorescence emission from cells was measured upon excitation with blue (488nm) laser. Post-acquisition data was analysed using Summit software V5.2.1.12465.

**Immunoblotting**

Antibodies used in the experiments are pERK1/2, pMEK1/2^S217/S221^, pACC^S79^, pAkt^S473^, pPEA15^S116^, tERK, tMEK, ACC, Akt, tPEA15, TFEB, myc tag, HA tag, cleaved PARP, LC3, IgG (Cell Signaling Technology), α Tubulin (Calbiochem), PHLPP2 (Abcam), Flag-tag (Sigma-Aldrich), LAMP2 (Abcam), p62 (Cell Signaling Technology) followed by HRP-tagged secondary antibody (Jackson ImmunoResearch). Chemiluminescence (using ECL substrate from Thermo Fisher Scientific) image was acquired by Syngene G-Box bio imaging system.

**Immunoprecipitation**

For co-immunoprecipitation, cells were lysed in IP-lysis buffer containing 25 mM Tris, 150 mM NaCl, 1 mM EDTA, 1% NP-40, 5% glycerol (pH 7.4). Lysates (1.5mg) were incubated with IgG control, anti-Flag, or anti-PEA15 antibody and 15 µL of protein-A sepharose beads for 12 hours at 4°C on end-on rocker. The immune complexes were washed with Nonidet P-40 lysis buffer (25 mM Tris, 150 mM NaCl, 1 mM EDTA, 1% NP-40, 5% glycerol; pH 7.4) for 5 times. Immunocomplexes were analysed by immunoblotting.

**Immunofluorescence**

Immunofluorescence on attached and suspension cells were done as described previously (Sundararaman, Amirtham et al. 2016). Briefly, cells in attached condition were fixed on dish by 4% PFA at room temperature for 10 minutes. Suspension cells were collected in 1.7 ml tubes and centrifuged at 3000 rpm for 3 min followed by 4% PFA fixation at room temperature for 10 minutes and spotted on coated glass slide. Permeabilization was carried out by 0.1 % triton X 100 for 15 minutes. Primary antibody diluted in PBST was added on cells and incubated overnight at 4°C or 2 hours at room temperature followed by incubation with fluorophore-tagged secondary antibody for 45 minutes at room temperature. Images were acquired using Olympus FV10i confocal laser scanning microscope after mounting the samples.

**Co-localization analysis**

The quantitative co-localization analysis of LAMP2 and GFP-LC3 was performed using Coloc-2 plugin in ImageJ program (NIH Image). For analysis, images of 30-40 cells were taken from multiple independent experiments (n=3). Each dot in the plots represents Pearson's correlation coefficient (PCC) value of LAMP2 and GFP-LC3 from a single cell. PCC is a statistical measure of the strength of the linear relationship between two data sets. Its value ranges from -1 to +1, with -1 representing negative co-relation, 0 no-co-relation and +1 represents a complete positive correlation.

**Real-Time qPCR analysis**

TRIzol reagent was used for isolation of total RNA. cDNA was prepared using random hexamer primers. Quantitative PCR was performed in Mastercycler RealPlex2 machine (Eppendorf) by using the KAPA SYBR FAST (Sigma-Aldrich). The primer sequence used in the study are: cFOS Forward primer: 5’AGTTCATCCTGGCAGCTCAC-3’ and cFOS Reverse primer: 5’ TGCTGCTGATGCTCTTGACA-3’.

**Immunohistochemistry (IHC)**

Day 7 lactating mammary gland was fixed in formalin, paraffin embedded and sectioned onto charged slides (Hisure Scientific). Slides were kept in air oven at 65°C overnight. Xylene was used to remove paraffin that is followed by rehydration in decreasing concentration of ethanol (100-70%). Tris-EDTA (pH 9) was used for antigen retrieval and 3% H_2_O_2_ solution for 20 minutes was used for neutralizing endogenous peroxidases. Primary antibody was incubated overnight at 4°C. Enhancer and secondary antibody were used as described by manufacturer’s instruction (Biogenex Supersensitive polymer HRP IHC detection kit). Diaminobenzidine (DAB) was used as substrate for peroxidase, while counterstain was done with haematoxylin. Images were acquired by IX71 Olympus inverted microscope.

**Experimental setup for *in vivo* metastasis studies**

EAC cells were cultured in suspension with RPMI media (with 10% FBS) for 48 hours in noble agar coated dish along with DMSO, as vehicle control, or PD98059. Viable cells (1x10^5^) were taken based on trypan blue exclusion staining and injected intraperitoneally in Swiss albino mice. Intraperitoneal injection of PD98059 was given on every alternate day till 15 days followed by dissection of lung after scarifying the animals. Tissues were fixed in paraformaldehyde and embedded in paraffin followed by thin sectioning. Lung metastasis was analysed on H&E stained slides of the tissue section. Images were taken using I X 71 Olympus inverted microscope.

Female Balb/c and Balb/c nude mice (6–8 weeks old) were procured from the Jackson Laboratory (Bar Harbor, ME, USA) and maintained in the Central Animal Facility of the Indian Institute of Science (IISc), Bengaluru, India. All animal procedures were performed in accordance with the guidelines of the Institutional Animal Ethics Committee (IAEC) and the Central Animal Facility, IISc, and were approved by the Institutional Animal Care and Use Committee (IACUC).IAEC Ethnic number for Female Balb/c (CAF\ETHICS\009\2023).

Female Balb/c mice (n = 5; 6–8 weeks old; 22–24 g) were injected via the tail vein with 0.15 mL phosphate-buffered saline (PBS) containing 4T1-eGFP cells (5 × 10⁶ cells/mL).Twenty four hours prior to injection, cells were seeded in soft agar-coated plates in a group of two to induce suspension-mediated stress and activate AMPK signaling. Among that one group was treated with PD98059 (20 µM; an ERK inhibitor), while a control group received an equivalent concentration of DMSO.

Two mice each were injected with cells from the control or PD98059‑treated groups, while one served as a no‑tumor control (NTC). Metastatic distribution and lung tumor growth were monitored by in vivo bioimaging. Fifteen days post‑cell injection, two mice received intraperitoneal PD98059 (10 mg/kg) and two received DMSO. After four weeks, mice were euthanized, lungs were excised, and metastatic nodules on the lung surface were enumerated to assess metastatic burden.

**Microarray data analysis**

MDA-MB-231 cells were cultured for 24 hours in attached or matrix-deprived condition and MDA-MB-231 cells stably expressing shAMPKα2 cultured in suspension condition. Microarray assay was done after harvesting cells at indicated time points. RNA isolation was performed using RNeasy minikit (Cat No. 74104; Qiagen). Cy3 labelling was done by Agilent's Quick-Amp labeling Kit (Cat. No. 5190-0442) followed by hybridization on Agilent's In situ Hybridization Kit (Cat No.5188-5242). The comparison was done between expression of same genes across test sample and control sample. The list of signature genes were selected from AmiGO Gene Ontology Consortium (<http://amigo.geneontology.org/amigo>), Profiler PCR Array list Qiagen, and further validated by KEGG (<http://www.genome.jp/kegg/pathway.html>) or PubMed (<https://www.ncbi.nlm.nih.gov/pubmed/>). The heat map of signature genes was created using online tool Morpheus ([https://software.broadinstitute.org/morpheus/#](https://software.broadinstitute.org/morpheus/)). Downregulated genes were color coded in “green” and upregulated genes in “red”. The enriched altered pathways in the suspension culture, were determined by Gene Set Enrichment Analyses (GSEA).

RNA sequencing data for single-CTC and cluster-CTCs were taken from Gene Expression Omnibus (GEO; <https://www.ncbi.nlm.nih.gov/geo/>), with GEO IDs GSE51827. The raw value was converted into log_2_ value. Box plots for the value of signature genes were plotted using GraphPad Prism 5.0 software.

**Mathematical modelling**

A mathematical model was constructed to depict the interaction between pAMPK, pERK and TFEB using a set of three coupled ODEs that consider the timescale separated kinetics of protein phosphorylation/dephosphorylation and protein production processes. Kinetic rate constants were estimated from previous studies and current work. A detailed description of the model is presented in the supplementary text. Nullclines and bifurcation plots were generated using MATLAB (Mathworks Inc.).

**Mathematical Modelling of pAMPK/pERK/TFEB system**

Our results show that at low pAMPK, ERK-PEA15 complex is driven more towards pERK-PEA15 and at high pAMPK, towards ERK-pPEA15. The pERK-PEA15 complex is responsible for a decrease in the stability of TFEB which is seen to increase AMPK activity by promoting phosphorylation of AMPK, completing the feedback loop (Figure 8E). The total ERK and total AMPK concentrations remain unchanged. Based on these observations, we construct a mathematical model with the following assumptions:

1. pERK destabilizes TFEB by increasing the degradation rate of TFEB
2. Phosphorylation and dephosphorylation reactions (of pAMPK and pERK) are much faster as compared to the destabilization effect of ERK on TFEB, such that any change in concentration of TFEB will immediately reflect in the concentration of pAMPK and pERK.
3. pAMPK depletes pERK-PEA15 (and hence reduces the inhibition effect on TFEB) by driving the dephosphorylation reaction. The effect is originally indirect (competitive phosphorylation of ERK-PEA15) but since our interest is only pERK-PEA15, we only consider depletion of it via pAMPK. This allows us to treat the complex as simply ERK.

Based on the above assumptions, the system can be formulated as follows:

$\frac{d\left[ TFEB \right]}{dt}=g$_TFEB_$-k$_TFEB_$*H$^s-^$\left( pERK, TFEB \right)$

$\left[ pERK \right]=Et*\frac{KeqERK}{1+KeqERK}*H$^s-^$\left( pAMPK, pERK \right)$

$\left[ pAMPK \right]=At*\frac{KeqAMPK}{1+KeqAMPK}*H$^s+^$(TFEB, pAMPK)$*$H$^s+^$(S, pAMPK)$

Where [TFEB], [pERK] and [pAMPK] denote the concentrations of TFEB, phosphorylated AMPK and phosphorylated ERK respectively. g_TFEB_ and k_TFEB_ denote the production and degradation rates of TFEB, KEqAMPK and KEqERK denote the equilibrium constants of phosphorylation of AMPK and ERK, Et and At represent the total ERK and total AMPK concentrations (constants) and $H$^s+/-^$\left( x,y \right)$ denotes the Hill shift function that results in the fold change in the rate due to x (Lu, Jolly et al. 2013). The hill shift function is defined as

$H$^s^$\left( x,y \right)=H$^-^$\left( x,y \right)+\lambda$_x,y_$*(1- H$^-^$(x,y))$

Where $H$^-^$\left( x,y \right)$ is the inhibitory Hill function defined as $H$^-^$\left( x,y \right)=\frac{1}{\left( x_{0y} \right)^{n_{x,y}}+\left( X \right)^{n_{x,y}}}$, $x_{0y}$is the threshold concentration of X for inhibition of Y, $n_{x,y}$ is the hill coefficient and *X* is the concentration of X. $\lambda$_x,y_ denote the fold-change in production/degradation rate of Y due to X. For $\lambda$_x,y_ >1, these functions are represented as $H$^s+^, and for $\lambda$_x,y_ <1, these functions are represented as $H$^s-^. *S* in the above equations represents an external signal affecting the phosphorylation of AMPK.

The degradation rate of TFEB was estimated based on a first order kinetics. From the stability study of TFEB, we estimated the half-life of TFEB to be ~ 15 hours and correspondingly the innate degradation rate as 0.052/hr (Figure S5E). The corresponding production rate was estimated by considering a *g/k* ratio (equilibrium concentration in the cell) of the order of ~10^5 molecules (Zeiler, Straube et al. 2012). The equilibrium constant of AMPK phosphorylation and rates of ERK phosphorylation and dephosphorylation were estimated from previous studies (Aoki, Yamada et al. 2011, Sonntag, Dalle Pezze et al. 2012). $\lambda$_TFEB,pAMPK_ and $\lambda$_pERK, TFEB_ were estimated for TFEB and AMPK based on the experimental data (Figure 7B, 6D) as 2.5 each. The hill coefficients were taken as 6 and the thresholds for each hill function were adjusted around the half-maximal concentration of the corresponding species.

The full list of parameters is given in the table below.

| g_TFEB_ | 500* |
| --- | --- |
| k_TEFB_ | 0.052** |
| KeqAMPK | 100000 |
| KeqERK | 100000 |
| $\lambda$_TFEB,pAMPK_ | 2.5 |
| $\lambda$_pAMPK,pERK_ | 0.1 |
| $\lambda$_pERK, TFEB_ | 2.5 |
| n_TFEB,pAMPK_ | 6 |
| n_pERK, TFEB_ | 6 |
| nae | 6 |
| t0a | 8000*** |
| a0e | 70000*** |
| e0t | 250000*** |
| At | 35000*** |
| Et | 470000*** |
| S | 100*** |
| s0a | 100*** |
| nsa | 1 |
| $\lambda$_S,AMPK_ | 2 |

Units:
* - molecules/hr
** - hr^-1^
*** - molecules

Nullclines for this three-dimensional system were plotted between two components of the system by assuming the third component to be in a quasi-steady state.

Phenotypic switching occurs via external perturbations that may or may not be correlated with the network under study. To demonstrate phenotypic switching upon perturbation, the system was simulated (using ode23s function of MATLAB) for a long time with intermittent stochastic perturbations. The perturbations were timed such that between two consecutive perturbations, system achieves a steady state. The external signal, S, was used as the bifurcation parameter to identify the bistable region for the system. Sensitivity analysis with respect to the bistable region was performed for the Hill coefficients and the total AMPK and ERK concentrations. While bistability is maintained upon variation in the Hill coefficients and the total ERK concentration, it vanishes for lower values of total AMPK. (Figure S8B). This is because the threshold of pAMPK to effect pERK is almost comparable with the maximum pAMPK levels produced at lower total AMPK levels. Bifurcation plots for sensitivity analysis were generated using MATCONT (Dhooge, Govaerts et al. 2003).

**Statistical Analysis**

GraphPad Prism V software was used to plot graph and analyse statistical significance of the data by using student’s t-test. Each experiment was performed at least thrice and one is being represented in the figures. All data are presented as mean ± standard error of the mean (SEM). *P* value below 0.05 was considered as statistically significant; ***represents *P* ≤0.001, **represents *P* ≤0.01 and *represents *P* ≤0.05.
